# RAMEN Unveils Clinical Variable Networks for COVID-19 Severity and Long COVID Using Absorbing Random Walks and Genetic Algorithms

**DOI:** 10.1101/2023.01.24.525413

**Authors:** Yiwei Xiong, Jingtao Wang, Xiaoxiao Shang, Tingting Chen, Douglas D. Fraser, Gregory Fonseca, Simon Rousseau, Jun Ding

## Abstract

The COVID-19 pandemic has significantly altered global socioeconomic structures and individual lives. Understanding the disease mechanisms and facilitating diagnosis requires comprehending the complex interplay among clinical factors like demographics, symptoms, comorbidities, treatments, lab results, complications, and other metrics, and their relation to outcomes such as disease severity and long term outcomes (*e*.*g*., post-COVID-19 condition/long COVID). Conventional correlational methods struggle with indirect and directional connections among these factors, while standard graphical methods like Bayesian networks are computationally demanding for extensive clinical variables. In response, we introduced RAMEN, a methodology that integrates Genetic Algorithms with random walks for efficient Bayesian network inference, designed to map the intricate relationships among clinical variables. Applying RAMEN to the Biobanque québécoise de la COVID-19 (BQC19) dataset, we identified critical markers for long COVID and varying disease severity. The Bayesian Network, corroborated by existing literature and supported through multi-omics analyses, highlights significant clinical variables linked to COVID-19 outcomes. RAMEN’s ability to accurately map these connections contributes substantially to developing early and effective diagnostics for severe COVID-19 and long COVID.

## Introduction

The outbreak of the COVID-19 pandemic has reshaped billions of lives across the globe and caused catastrophic socioeconomic losses in the past years[1, 2]. Despite the impact that COVID-19 has brought to humanity, a comprehensive understanding of COVID-19 disease mechanisms remains elusive, which substantially limits the diagnostics and therapeutics for COVID-19. For instance, it is known that different COVID-19 patients develop distinct symptoms and clinical outcomes[3]. However, it remains unclear why some people tend to develop more severe COVID-19 outcomes while some barely show any symptoms. This lack of understanding restricts the early clinical diagnosis and interventions for the most vulnerable COVID-19 victims[4, 5], which leads to undoing suffering. Moreover, although most COVID-19 victims can recover from the disease, approximately 10-30% of them develop long-term symptoms (termed as long COVID or post-COVID-19 conditions)[4], which has a tremendous socioeconomic impact. Affected individuals endure both physical and psychological distress[5]. The loss of work capacity among many sufferers has notably hindered economic productivity. The fast-accumulating COVID-19 datasets present unprecedented opportunities to derive a deeper understanding of the disease mechanisms underlying COVID-19, which will drive the discovery of novel diagnostic markers and therapeutic targets. Multiple initiatives across the globe are profiling clinical variables and genomics sequencing associated with COVID-19 patients[6, 7]. In Quebec, Canada, the Biobanque québécoise de la COVID-19 (BQC19)[8] has collected clinical data for over six thousand COVID-19 patients, along with data from several other modalities including proteomics and transcriptomics on a substantial subset. In Ontario, Canada, investigators at the Lawson Health Research Institute have also collected COVID-19 patient clinical information data [9]. These valuable data resources provide us with the opportunity to develop novel computational methods for better understanding the disease, and coming up with potential diagnosis methods and treatments,

With the ever-increasing availability of COVID-19 datasets, many studies interrogated the complex relationships between various clinical variables and COVID-19 severity levels [10–13] using simple statistical methods (e.g., correlation[13] and mutual information [14]). Existing methodologies excel at mapping direct relationships between clinical variables and outcomes, delineating these associations as edges within a network. However, while standard statistical tools like Pearson correlation and mutual information are proficient at inferring associations between variables, they do not elucidate the directionality of these relationships. These methods might identify direct connections between pairs of variables, yet they often overlook the indirect relationships crucial in complex networks composed of hundreds of variables. Moreover, statistical approaches focused on mutual information or correlation primarily examine pairwise relationships, leaving the complex interactions among multiple variables largely unaddressed. Additionally, although direct correlational analysis can effectively highlight associations between two variables based on direct correlation tests, it does not necessarily imply that these associations are relevant or predictive of the disease outcome. In essence, the relationships identified through correlation analysis may not be directly connected to the disease outcome, revealing a significant limitation in the application of such analyses for comprehensively understanding disease mechanisms and their impacts.

On the other hand, Bayesian Network (BN)[15–17], a probabilistic graphical model, can address the above limitations of simple statistical approaches, enabling the inference of clinical variables that could indicate COVID-19 outcomes (e.g., COVID-19 severity or long COVID). Previously, BNs have shown success in many application scenarios and outperformed physicians in disease diagnostics[18, 19]. For example, a BN-based diagnosis not only achieved state-of-the-art overall performance for neurodegenerative diseases compared to a list of other methods [20–25] but also provided very good interpretability. However, building a BN (particularly structure learning) for hundreds and thousands of clinical variables is challenging and computationally expensive. This task is complicated by the huge search space of possible solutions (In fact, the problem is NP-hard[26]). In practice, the BN structure learning is often regularized by prior knowledge (e.g., known constraints) to reduce the search space[27, 28]. Unfortunately, this is not an option for COVID-19 since our current understanding of this relatively new disease remains very limited[29]. Additionally, since our major aim is to find new clinical variables that influence COVID-19 outcomes (COVID-19 severity or long COVID) either directly or via another clinical variable, relying on prior knowledge unavoidably introduces bias.

To address the above limitations and fill the gap, here we introduce RAMEN (Random walk and genetic AlgorithM-based nEtwork iNference) that glues random walk and Genetic Algorithm to infer a Bayesian network representing the relationships between clinical variables in COVID-19 (particularly between clinical variables and COVID-19 outcomes). The random walk is employed to rank and select the most relevant variables and connections to COVID-19 outcomes to reduce the network complexity.

A significant aspect of our methodology is the incorporation of a terminal absorbing node, symbolizing the disease outcome of interest, such as COVID-19 severity, within our clinical variable network. In the process of conducting a random walk, all paths culminate at this terminal absorbing node. This approach effectively assigns a “direction” towards the disease outcome for the random walk, thereby facilitating the identification of network edges (associations) that have a direct link to the disease outcome (absorbing node). This innovative feature enhances the capability to discern which associations are most relevant to understanding and predicting the disease outcome. Following the preliminary network reconstruction through the absorbing random walk process, we employ a Genetic Algorithm to refine and identify an optimized network structure. This optimized structure is more accurately aligned with the observed clinical variables, ensuring a precise representation of the relationships and interactions within the dataset. After these two stages, RAMEN outputs a BN that models the complex relationship between the COVID-19 outcome variable (*e*.*g*., severity and long-COVID) and other variables that are directly or indirectly connected to it. To examine the performance of RAMEN, we applied the method on three different COVID-19 cohorts from the BQC19 project and Lawson Health Research Institute, examined the resulting network with multi-omics measurements and computational simulations, and compared it with other methods. We show the resulting networks capture important COVID-19 outcome indicators that can be validated via multi-omics, simulation, or literature. Moreover, RAMEN demonstrated superior performance over simple statistical methods by finding more relevant variables and indirect variables that cannot be found using simple statistical methods such as mutual information and Pearson correlation.

Our model’s ability to predict early indicators of COVID-19 outcomes, such as severity and long COVID, significantly enhances early diagnostic development for patients at risk of severe symptoms or long COVID. This advancement enables personalized and timely diagnosis, streamlining targeted treatment strategies. Such precision in diagnosis and treatment not only improves patient outcomes but also bolsters our fight against the pandemic by allowing for more efficient resource allocation and reducing the spread and impact of the virus on individuals and healthcare systems. Furthermore, this model has the potential to be generalized for analyzing clinical variable networks in diseases beyond COVID-19, provided that similar records of clinical variables are available. This enhances its utility across a wider spectrum of medical research.

## Results

### Overview of RAMEN

The RAMEN method comprises two phases: first, RAMEN applies absorbing random walks to select the most relevant variables to the COVID-19 outcomes (severity or long COVID) and builds a draft candidate network. Second, RAMEN employs a Genetic Algorithm to search the optimized network that represents the relationships between different clinical variables based on the draft network from the random walks (Fig. 1).

**Fig. 1.**
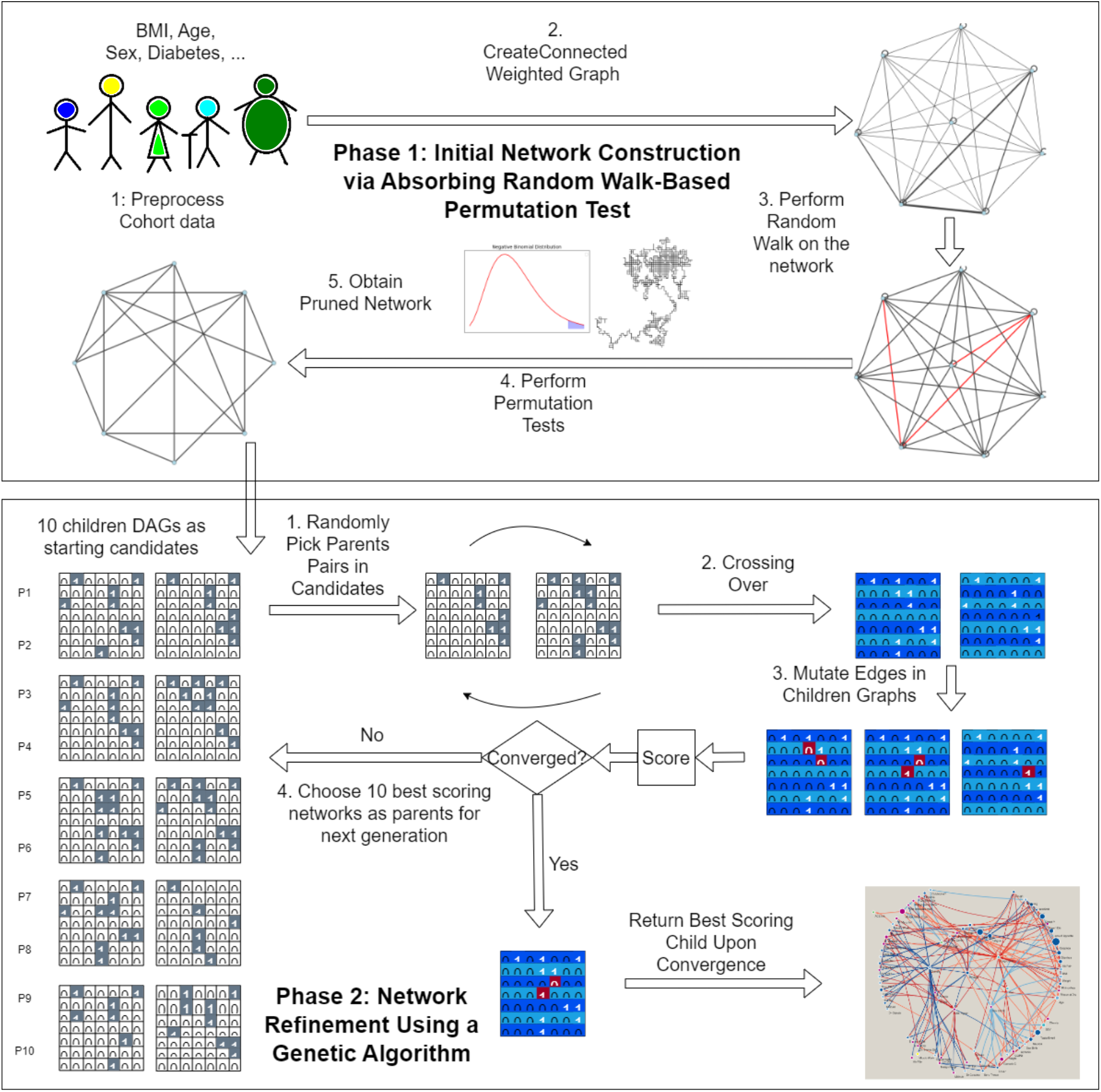
Overview of the RAMEN Methodology. The RAMEN approach constructs Bayesian networks from clinical data through a sequential two-phase process. **Phase 1: Establishing the Initial Network with Absorbing Random Walk-Based Permutation Test**. Beginning with preprocessed clinical data, this stage implements a permutation test via a random walk strategy across a comprehensive network of all included variables, where nodes symbolize variables and edge weights indicate the mutual information among variable pairs. The process identifies stronger variable connections by tracking the frequency of edge traversal in successful random walks (ending at the target node). Edges with significantly higher traversal frequencies, as established through permutation testing, lay the groundwork for the network, preparing it for further enhancement. **Phase 2: Enhancing the Network with a Genetic Algorithm**. This stage further refines the Bayesian network structure. Starting with a set of initial network configurations derived from the early framework, the Genetic Algorithm applies crossover (merging two configurations) and mutation (applying random changes) to evolve these structures. Each cycle assesses the network structures against a specific scoring function, prioritizing those with superior scores for subsequent iterations. This cycle of refinement, through modification, assessment, and selection, persists until a stable score is achieved, culminating in an optimized network structure.

For the random walk phase 1, we first initialize a full network composing all variables as nodes, and then compute edge weights as the mutual information between the node pairs. The clinical variable of interest (often disease outcome) will be regarded as the absorbing node, for which only incoming edges are allowed. All other nodes in the graph will be regarded as non-terminal. We then perform random walks starting from each non-terminal variable (all variables other than the COVID-19 outcomes) for *N* random steps. At each node, the random walk uses the normalized mutual information stored in outgoing edges as transition probabilities. A random walk will terminate successfully if and only if it reaches the absorbing (terminal) node within *N* steps. Edge visits of successful random walks will be recorded. Intuitively, edges with more visits are the ones with stronger connections and are relevant to the COVID-19 outcome node. Subsequently, to find the edges with significantly more visits, we perform a permutation test. Specifically, we run another version of random walk on a network with permuted edge weights and use the edge visits in this network as the random background. Finally, we fit a Negative Binomial distribution to the random background (see Supplementary Fig. S1 for the histogram of edge visit count) and select edges with significantly more visits (q-value ≤ 0.05) to form the initial network skeleton.

Utilizing the Genetic Algorithm (GA), our method refines the Bayesian network structure by starting with an initial network skeleton and generating a diverse pool of initial network candidates in phase 2. These candidates are created by randomly assigning directions to the bidirectional edges in the skeleton. The evolutionary process involves performing crossover between candidate pairs to produce combined networks (offspring), introducing mutations to these offspring to create variability, and then scoring all candidates—parents, offspring, and mutated offspring—using an Entropy-based scoring function. The highest-scoring networks are selected for the next iteration. This cycle of crossover, mutation, evaluation, and selection is iterated until the scoring converges, at which point the final network is obtained. This approach systematically explores and refines potential network structures, ensuring the development of an optimized Bayesian network that accurately models the clinical data.

### Building COVID-19 severity network using the BQC19 hospitalized patient dataset

In our study, the RAMEN methodology was applied to the hospitalized patient cohort data from BQC19, resulting in the development of a Bayesian Network (BN). This BN delineates the complex interplay among clinical variables and their association with COVID-19 severity (see Fig. 2a). Our analysis encompassed 2,018 hospitalized patients, with the dataset including 297 clinical variables post-cleaning. The severity of COVID-19 within this cohort was classified into three categories: “not infected or mild,” “moderate,” and “severe or deceased.”

**Fig. 2.**
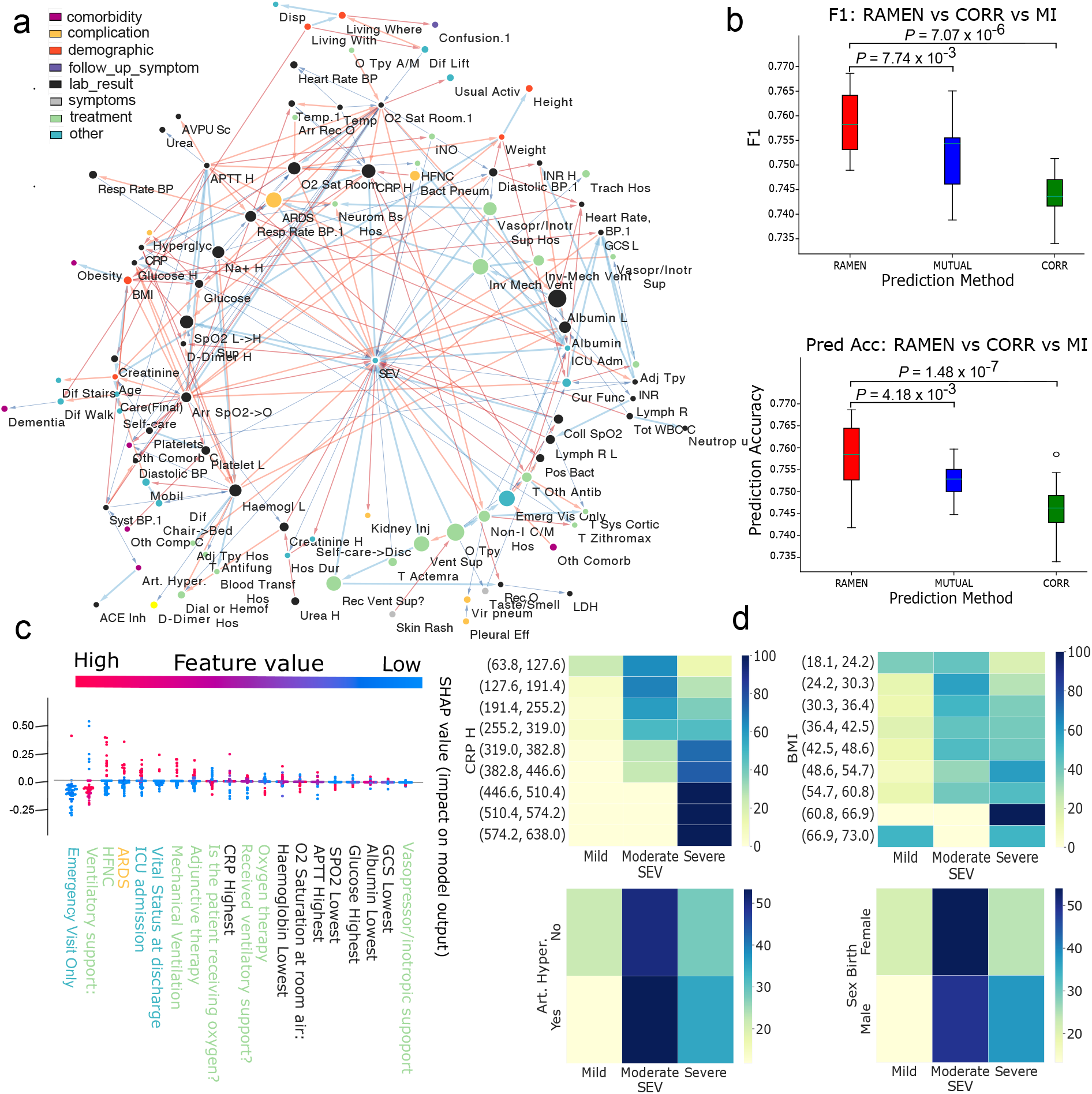
RAMEN unveils indicators of COVID-19 severity in BQC19 hospitalized patient data. **a**, A streamlined network showcasing 231 of the most significant connections identified by RAMEN, indicative of COVID-19 severity. The full names of the variables are in Supplementary Table S1. The color and thickness of edges signify the connection strength (blue for weaker, red for stronger) based on mutual information metrics. Nodes are colored according to categories of clinical variables, with their size reflecting the strength of their correlation with COVID-19 severity. **b**, Comparison of F1 scores for predicting COVID-19 severity, contrasting RAMEN-identified indicators against those identified through mutual information (MUTUAL) and Pearson correlation (CORR) methods, with predictions made by Support Vector Machines (SVMs). A higher F1 score suggests a greater relevance of the identified variables for severity prediction. **c**, Analysis of SHapley Additive exPlanations (SHAP) values, elucidating the significance of clinical variables pinpointed by RAMEN in SVM-based predictions. These values depict the impact of variables on the model’s prediction as either contributing towards a positive or negative outcome, with a consistent color scheme across the x-axis reflecting dependable predictors. **d**, Heatmaps illustrating the conditional distribution of COVID-19 severity levels (SEV) across the values of direct indicator variables, where the heatmap colors represent the proportion of patients within each severity category for given indicator values. This visual representation aids in understanding the correlation between specific clinical indicators and severity outcomes.

The clinical variable network, as inferred in relation to COVID-19 severity, aligns with findings reported in the existing literature. Among the variables identified, several key examples linked to “COVID-19 severity” include “Sex”[11, 30], “Age”[11, 30], “BMI”[31, 32], “Arterial Hypertension”[33], “ALT”[34], “C-reactive protein (CRP) (highest value)”[35], and “Albumin (lowest value)”[36]. These variables represent just a sample of the broader network, illustrating the diverse factors impacting COVID-19 severity as observed in our study.

To demonstrate the superior capability of RAMEN in identifying relevant indicators for COVID-19 outcomes compared to conventional statistical methods, we conducted benchmark analysis focused on predicting COVID-19 severity based on the early record of clinical variables. This involved training a Support Vector Machine (SVM) classifier using indicator variables from the COVID-19 severity network established by RAMEN, and two additional SVMs, each utilizing the top variables identified by mutual information and Pearson correlation methods, respectively. The effectiveness of these indicators was assessed based on the predictive performance of the SVMs. As depicted in Fig. 2b, the F1 scores and accuracy of the three SVMs, corresponding to the variables identified by each method, are presented on the x-axis. The p-values from the Wilcoxon test, comparing RAMEN with the other methods, indicate significantly higher performance for predictions based on RAMEN-identified variables. This result underscores RAMEN’s ability to uncover more pertinent COVID-19 outcome indicators than traditional statistical methods. The developed classifier can also be used to predict the outcome of the disease (such as the severity of COVID-19) based on the early clinical variable records from the first month of patient care.

To assess the effectiveness of the indicators identified by RAMEN, we visualized the SHapley Additive exPlanations (SHAP) values in Fig. 2c. This visualization details how each indicator contributes to the SVM’s positive or negative predictions, with the y-axis representing the contribution magnitude and the x-axis listing the RAMEN-identified indicator variables. The plot reveals a consistent pattern of value distribution (as indicated by colors) across both sides of the x-axis, with many variables situated significantly away from the axis. This indicates that these indicators are reliable predictors of outcomes, further affirming the effectiveness of the indicators identified by RAMEN.

The association between identified indicators and COVID-19 severity is further elucidated through heatmaps, as shown in Fig. 2d. These heatmaps detail the relationship between COVID-19 severity levels and the pertinent indicators identified by RAMEN. Each heatmap illustrates the variation in the percentage of patients across different severity levels in relation to the values of variables directly linked to COVID-19 severity. Generally, for variables that are strongly connected to COVID-19 severity within our reconstructed network, there is a significant shift in the distribution of severity levels corresponding to the values of these indicator variables. Conversely, for variables deemed irrelevant, the severity distribution remains largely unaffected by changes in these variables. The four heatmaps showcased underscore RAMEN’s efficacy in pinpointing highly relevant indicators of COVID-19 severity.

### Systematic validation of COVID-19 severity indicators identified by RAMEN using BQC19 multi-omics data

To validate the reliability of the COVID-19 severity indicators identified by RAMEN, we utilized the BQC19 multi-omics dataset. Our validation approach included a comparison of differentially expressed (DE) genes and proteins associated with each severity indicator against those associated with various levels of COVID-19 severity. This process involved examining the overlap between DE genes (from RNA sequencing data) or proteins (identified through SomaScan 5K array) related to the indicators and those distinguishing between mild and severe COVID-19 cases. The heatmaps depicted in Fig. 3a demonstrate a significant overlap of DE genes between the indicators and COVID-19 severity, revealing distinct expression patterns across the range of indicator values. These findings suggest that a common set of genes may be involved in linking these indicators to COVID-19 severity, indicating underlying biological pathways. Furthermore, the BQC19 multi-omics dataset provides insights into the biological mechanisms potentially governing these relationships. Supplementary Fig. S2 shows an additional example between “ARDS” and Severity. Pathway enrichment analysis of the “common” DE genes associated with COVID-19 severity and its primary indicators, shown in Fig. 3b, identified significant pathways including “Neutrophil degradation [37, 38],”Innate Immune System[39],” “Antimicrobial peptides[40],” and “Heme Signaling[41].” These pathways are implicated in modulating COVID-19 severity, suggesting mechanisms through which the indicators may influence disease severity. Severe COVID-19 is often characterized by acute respiratory distress syndrome (ARDS) associated with abnormal coagulation [42, 43]. Moreover, several studies have pointed to neutrophilia, release of their granules and neutrophil extracellular traps (NETs) as key pathological features of thrombotic complications driven by the immune system (also called immuno-thrombosis) in severe COVID-19 [3, 44–54], linking “ARDS”, “Neutrophil degradation”, “Innate Immune System” and “Heme Signaling”. Taken together, the pathway enrichment performed using the severity indicators identified by RAMEN is congruent with the existing literature, supporting the validity and reliability of RAMEN. We have also done this analysis using SomaScan data, which is illustrated in Supplementary Fig. S3, S4, and S5.

**Fig. 3.**
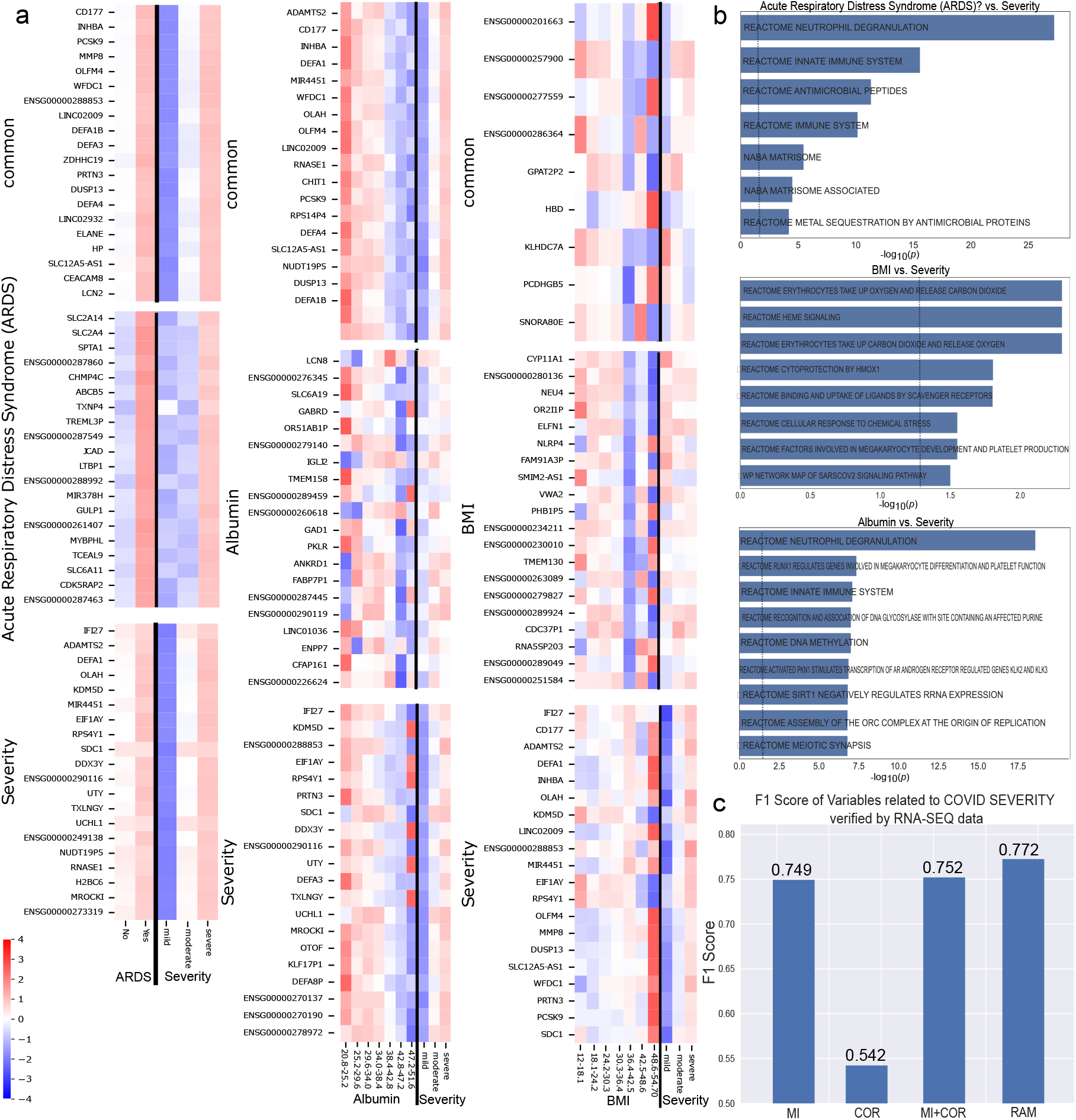
Support for the COVID Severity network edges from the RNA-seq data. **a**, Analysis of gene expression across three groups of differentially expressed (DE) genes linked to example nodes “ARDS”, “Albumin”, and “BMI” that directly connected to COVID severity. For example, with “Albumin”, we first pinpoint DE genes associated with Albumin variability (i.e., genes with expression changes in patients with varying Albumin levels, denoted as *G*_1_). Next, we identify DE genes linked to COVID severity (*G*_2_). The “Common” group in the figure represents DE genes common to both sets (*G*_1_ ∩ *G*_2_); the “Albumin” group illustrates DE genes exclusive to the Albumin variable (*G*_1_ ∩ *¬G*_2_); and the “Severity” group shows DE genes unique to COVID-19 severity (*G*_2_ ∩ *¬G*_1_). **b**, Identification of the top enriched pathways for each variable based on their common DE genes with the severity variable (the “common” group). The x-axis shows the negative log 10 of FDR-corrected p-values. **c**, Validation of COVID-19 severity indicators using various methodologies. Each method on the x-axis (MI: mutual information; RAM: RAMEN; COR: Pearson correlation) classifies variables into indicators or non-indicators, with RNA-seq data providing the basis for ground truth. A variable is considered an indicator if its DE genes significantly overlap with those associated with COVID-19 severity, assessed via a hypergeometric test. The classification performance of each method is quantified using the F1 Score from verifying the variables found by each method against the ground truth.

In addition, we performed a systematic benchmarking of RAMEN against other statistical methods, such as mutual information and Pearson correlation, to assess its effectiveness in identifying severity indicators (Fig. 3c). In this benchmarking exercise, RNA-seq data served as the basis for establishing a definitive classification of ground truth. The hypergeometric test was applied to assess the congruence between differentially expressed (DE) genes from selected indicators and those associated with COVID-19 severity, using p-values to determine statistical significance (see Methods). Clinical variables demonstrating a significant overlap of DE genes with COVID-19 severity were acknowledged as true indicators of severity for benchmarking purposes. The effectiveness of each method was assessed through the F1 score. According to Fig. 3c, correlation exhibited the lowest F1 score by a considerable margin, with Mutual Information showing significant improvement, yet still trailing behind RAMEN. The combination of correlation and mutual information was also evaluated, resulting in a marginally improved F1 score, though still not surpassing RAMEN. These results underscore RAMEN’s distinctive capacity to identify severity indicators that traditional statistical methods may fail to detect.

We have also done the same benchmarking using the SomaScan data, which is shown in Supplementary Fig. S6. Similarly, correlation exhibited the lowest F1 score by a considerable margin, with mutual information showing significant improvement. The combination of correlation and mutual information resulted in an improvement from the two methods individually. However, RAMEN still demonstrated a better score than all 3 methods.

### RAMEN reconstructs long COVID network from BQC19 outpatient data

We also applied RAMEN to the outpatient COVID-19 cohort to study long COVID. The outpatient dataset complements the hospitalized patients, having 1,440 patients and their 84 clinical variables. Whether or not an outpatient has long COVID is classified according to a commonly used criterion: the presence of at least one symptom at three months that persists and cannot be explained by pre-existing conditions[55]. In this cohort, based on the definition given above, long COVID frequency is 36.5% (526 out of 1440). It is important to note that this is not a random populational cohort, explaining the higher frequency. One of our objectives is to identify critical indicators for long COVID that may assist in the early diagnosis. For this study, we only used clinical variables profiled within one month after COVID-19 infection to make sure that the indicators that we found were meaningful for early diagnosis. RAMEN reconstructed a long COVID BN modeling the relationships between the clinical variables, with 36 indicators directly linked to long COVID (Fig. 4a). Many known critical variables for long COVID are captured, such as “Age”[56], “Chest”pain”[5], “Joint Pain”[57], “Runny nose”[57], and “Shortness of breath”[5]. On the other hand, irrelevant variables for long COVID, such as “chronic kidney disease”, are not shown in the network. This variable, to our best knowledge, is not reported to indicate a higher risk of long COVID. As discussed in one recent study[58], an in-depth assessment of kidney outcomes in the post-acute phase of COVID-19 infection is not yet available.

**Fig. 4.**
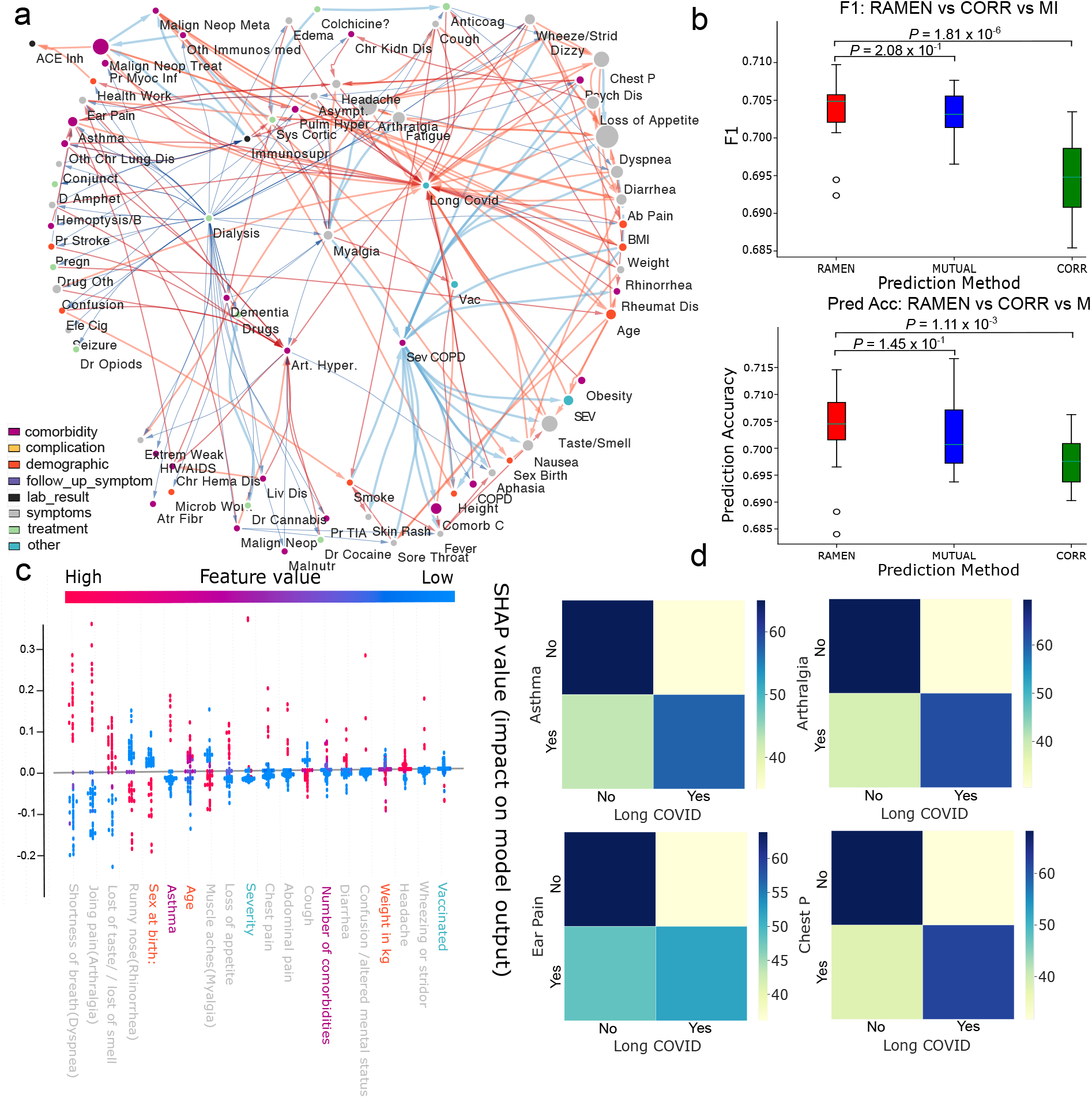
RAMEN identifies long COVID indicators from BQC19 outpatient data. **a**, Long COVID Network constructed using RAMEN. The full names of the variables are in Supplementary Table S1. Edge colors indicate connection strength (blue for low, red for high), node colors represent types of clinical variables, and node sizes show their relevance to long COVID. **b**, F1 scores for long COVID prediction using SVMs trained on indicators identified by RAMEN, mutual information (MUTUAL), and Pearson correlation (CORR). Error bars are plotted based on 20 5-fold cross-validations and p-values are from the one-sided Mann-Whitney U test examining if one method generates significantly better results. **c**, SHAP values showing the quality of long COVID indicators identified by RAMEN. **d**, Heatmaps showing the conditional distribution of long COVID variable, conditioned on direct indicators’ values.

While we have done the multi-omics analysis for long COVID as well, all the patients studied for long COVID are outpatients, meaning that we do not have enough multi-omics measurements for the patients to conduct a robust analysis. However, We have shown sample RNA-seq heatmaps, SomaScan heatmaps, and pathway enrichment analysis in Supplementary Fig. S7, S8, S9, S10, S11, and S12.

We demonstrate that RAMEN identifies early indicators (early records of clinical variables within the 1st month of patient care) for long COVID with enhanced precision, evidenced by the improved performance metrics (F1 score and accuracy) of an SVM classifier trained using these indicators, compared to SVM classifiers trained with indicators from other statistical methods. This indicates that RAMEN’s methodology in identifying indicators specifically for long COVID effectively enhances the accuracy of predicting the disease’s outcome when utilizing the SVM classifier (Fig. 4b). P-values from the onesided Mann-Whitney U test show that RAMEN results in significantly higher accuracy and F1 score than Pearson correlation. In addition to the disease outcome prediction performance, in Fig. 4c, we also show the high feature quality of indicators found by RAMEN with SHAP values. Most indicators found are very strong early predictors of long COVID. Finally, Figure 4d illustrates the influence of specific indicators on the long COVID variable by demonstrating the shifts in the distributions of long COVID values based on the indicator values. The four variables presented are all positively associated with long COVID; for instance, the presence of “Chest Pain” (ChestP) suggests an increased risk of long COVID.

To rigorously analyze the association between the outcome variable (long COVID) and the primary indicators, we employed Pearson’s chi-square tests. The outcomes, elaborated in the Supplementary Table S2, reveal a pronounced dependency between the top indicators identified by the RAMEN network (here we showed 9) and the long COVID outcome, underscoring the robustness of these indicators in relation to the disease. To quantify the distinctiveness of these findings from potential random occurrences, we estimated the background probability that a randomly chosen variable might significantly affect the disease outcome. This involved selecting sets of 10 clinical variables from a pool excluding those identified by the RAMEN network and assessing their genomic data support. Repeating this process 100 times yielded an average of merely 1.78 out of the 10 randomly chosen variables being substantiated (equating to a background probability of 0.0178). This step was crucial to establish a benchmark for comparison, demonstrating the specificity with which the RAMEN network’s top indicators correlate with the long COVID outcome, as opposed to the general expectancy of significance from a random selection of variables. The significant p-value from the binomial test (3.193 * 10^−8^) further confirms the non-randomness of the RAMEN network’s findings. This stark contrast highlights the precision and relevance of the RAMEN network in pinpointing critical indicators that have a meaningful dependency relationship with the disease outcome, significantly surpassing random expectation levels.

To evaluate the robustness and consistency of RAMEN, we applied it to an independent long COVID dataset from the Lawson Health Research Institute[9], which comprises 66 patients and 27 clinical variables, to ascertain if the outcomes were consistent with our previous long COVID study using BQC19 data. The results, depicted in Supplementary Fig. S13a, illustrate the network’s output using the Lawson Health Research Institute data. This dataset contains 8 variables that are common with the BQC19 dataset; of these, only 4 were identified by RAMEN as indicators of long COVID in the BQC19 dataset. These four indicators—”chest pain,” “anosmia/ageusia,” “dyspnea,” and “headache”—were also identified by RAMEN as indicators of long COVID in the independent dataset and are highlighted in red. The identification of these indicators is supported by previous research, indicating their relevance to long COVID [5, 59–61]. Additional variables discovered in this dataset provide further insights into the pathology of long COVID. The heatmaps in Supplementary Fig. S13b demonstrate the significant impact of these indicators on the long COVID variable, with the conditional distribution of the long COVID variable showing dramatic shifts in response to these indicators.

### RAMEN unveils variable relationships unreachable by mutual information or Pearson correlation for COVID-19 outcomes

To show that RAMEN can find extra information beyond simple statistical methods, especially finding network edges that cannot be reached via other methods, we tested RAMEN against Pearson correlation and mutual information. Fig. 5a and 5b show that RAMEN was able to find many edges that the other two methods were not able to find. We can see that it is especially the case in Fig. 5a, with 58.1% of the network edges being RAMEN unique. The full severity network has 79.8% being RAMEN unique. However, since Fig. 5b only shows 25% of top edges based on mutual information, there are only 22.5% of edges that are RAMEN unique.

**Fig. 5.**
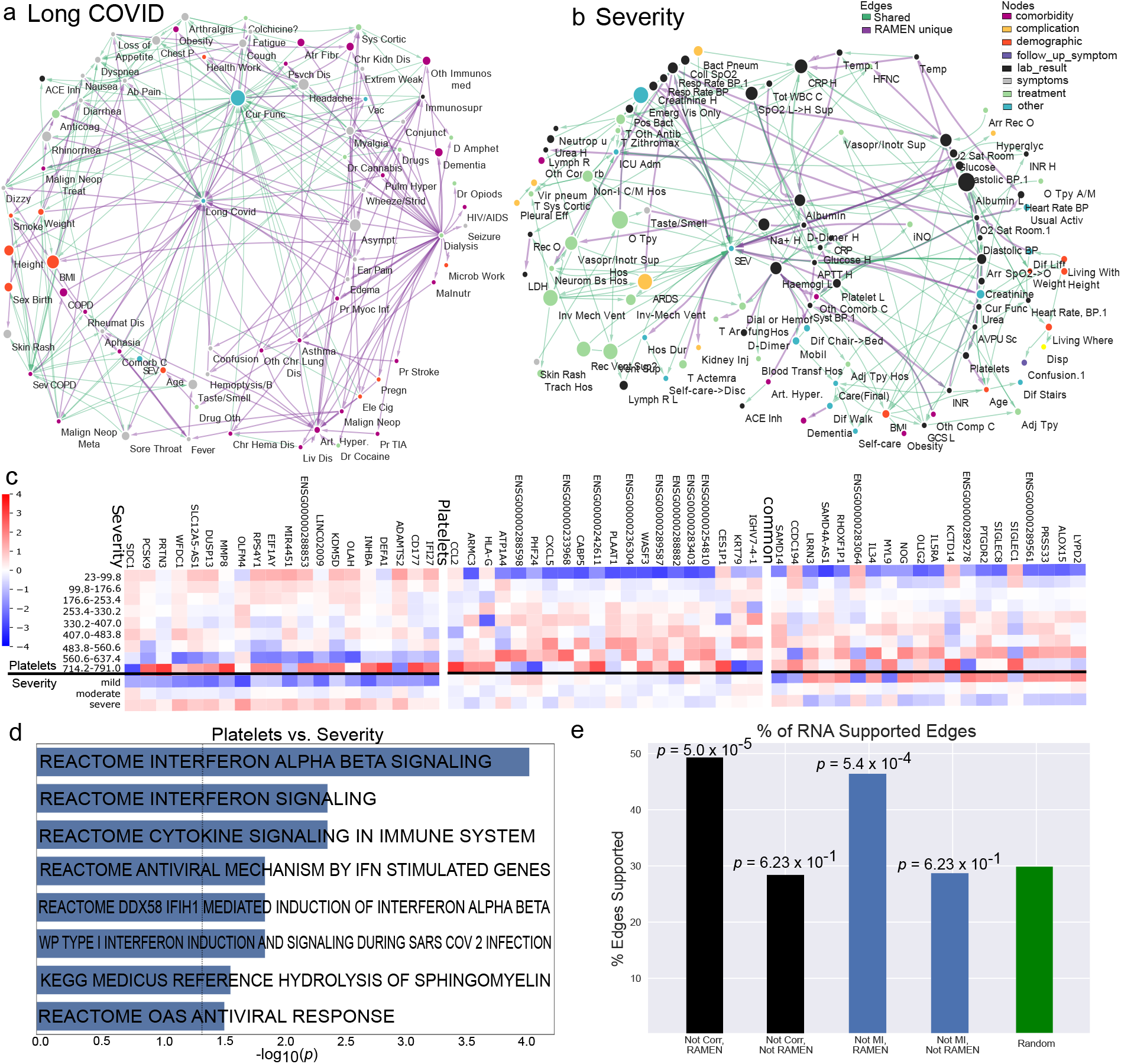
RAMEN identifies effective indicator variables that cannot be found using mutual information or Pearson correlation. **a**, The Long COVID network where purple edges represent connections significant only to RAMEN, and green edges are also identified by mutual information or correlation. **b**, Similar network for COVID-19 severity, with purple indicating edges found exclusively by RAMEN, and green representing those also found by mutual information or correlation. The full names of the variables are in Supplementary Table S1. **c**, Heatmaps visualizing DE genes associated with “Platelets” and “COVID-19 severity”. The three groups of DE genes correspond to the unique DE genes of the two variables and common DE genes. **d**, Pathway enrichment based on the common DE genes in **c. e**, A bar plot demonstrating RAMEN’s ability to detect disease-relevant edges missed by Pearson correlation and mutual information. Using RNA-seq data as ground truth (see the Methods section* for details), among all the edges that cannot be found using Pearson correlation, the column “Not Corr, RAMEN” shows the percentage of disease-outcome-relevant edges found by RAMEN. “Not Corr, Not RAMEN” shows those that also cannot be found using RAMEN. Likewise, “Not MI, RAMEN” corresponds to the percentage of true edges missed by mutual information but found by RAMEN, and “Not MI, Not RAMEN” are the ones that are not found by both. “Random” is the performance of random selecting edges. The p-values of the binomial tests indicate that RAMEN is accurate in finding edges missed by other methods. This suggests that RAMEN has additional power in detecting disease-relevant edges compared to Pearson correlation and mutual information.

In Fig. 5a, in the Long COVID network analysis, RAMEN identified 151 associations (edges) that were not detected by traditional correlational methods. By employing an absorbing random-walk approach integrated with a genetic algorithm, RAMEN was capable of identifying numerous associations that, while not demonstrating a strong direct correlation, play a significant role in influencing the random walk towards the absorbing node (disease outcome). This includes indicators of long COVID, such as “BMI:— Long Covid[62]”. Also, it includes edges such as “COPD (emphysema, chronic bronchitis) ?— Long Covid”, which is known to affect COVID severity[63]. This opens discussions to whether COPD can also be an indicator of long COVID. However, this could also be explained by the overlap of respiratory of symptoms between COPD and respiratory manifestations of long COVID, making it difficult to distinguish the etiology of the symptoms between these two conditions. Whether or not this link is a true biological association remains to be determined. Nevertheless, it demonstrates the usefulness of RAMEN in critically examining the data and the associations identified, promoting a better critical understating of pathogenesis. RAMEN can also find relationships between variables that are not directly related to long COVID, such as “Age at recruitement:— Loss of appetite ?[64]”. Furthermore, indirect connections, such as “BMI:— Rheumatologic disease ?— Long Covid”[65, 66], allow us to infer the relationships between the clinical variables in the case of long COVID. While correlational methods are able to find the relation between weight and BMI significant, it is not able to suggest the rest of the sequence of relationships. In this case, we know that weight is generally a risk factor for health, hence will likely be a risk factor for long COVID. However, through RAMEN, we are able to suggest the other clinical variables that weight might provoke which might be more precise risk factors and indicators of long COVID. Accordingly, Mendelian randomization studies showed that BMI can be causally linked to rheumatoid arthritis [67, 68]. The inflammatory component of rheumatologic disease driven by BMI could interact with long-term manifestations of SARS-CoV-2 infections, generating testable hypotheses based on RAMEN-identified vectors that may help us better understand long COVID in different subgroups.

In Fig. 5b, in the compact COVID severity network, correlational methods were not able to find 52 edges that RAMEN has found (They were not able to find 739 in the complete network). To give a few,”Creatinine (HIGHEST value)— COVID Severity[69]”, “Respiratory rate (associated with BP above):— COVID Severity[70]”, “COVID Severity— Acute kidney injury?[71]”. Acute Kidney Injury is a condition that often develops in patients affected with COVID-19, not only RAMEN was able to capture it while naive methods cannot, RAMEN was also able to capture the correct edge direction. Another edge that only RAMEN was able to capture that is worth pointing out is “Platelet (LOWEST value)— COVID Severity [72]”. Platelets occupy a major function in the immune system and have been found to be an indicator of COVID severity. Naive methods not being able to find such edges show that it is lacking compared to RAMEN in identifying risk factors that can worsen COVID severity.

In Figure 5c, similar to the previous figures, we demonstrate the overlap of differentially expressed (DE) genes between “Platelets” and “COVID severity”, revealing distinct expression patterns associated with the variables’ differing values. This analysis suggests the biological mechanisms underlying the predicted relationship between platelet levels and COVID-19 outcomes. In Figure 5d, a pathway enrichment analysis of the “common” DE genes related to both Platelets and COVID severity has identified several significant pathways. These pathways include “Reactome interferon alpha beta signaling” [73], “Reactome interferon signaling”, “Reactome cytokine signaling in the immune system”[74], “Reactome antiviral mechanism by IFN-stimulated genes” [75], “Reactome DDX58 IFIH1-mediated induction of interferon alpha beta” [76], “WP Type I interferon induction and signaling during SARS-CoV-2 Infection”[77], “Kegg Medicus reference hydrolysis of sphingomyelin”[78], and “Reactome OAS antiviral response”[79]. Type I interferons are established immunological mediators of COVID-19 severity [80–82]. Interestingly, platelets are key regulators of coagulation, a process that can be severely disrupted in severe COVID-19 leading to life-threatening conditions[83]. These identified pathways provide a biological context for the edges predicted by the analysis, confirming their relevance to the severity of COVID-19.

We further utilized RNA-seq data to validate the edges identified by RAMEN, particularly those missed by traditional correlational methods (MI and correlation). Our validation method hinges on the principle that two variables are considered connected if there is significant overlap in their differentially expressed (DE) genes, as outlined in the Methods section. Notably, RAMEN demonstrated a significantly higher capability to identify genomics-supported edges missed by correlation methods. This contrast becomes even more pronounced when examining edges that both RAMEN and the correlational methods failed to predict; these missed edges do not exhibit any enrichment in the number of supported edges compared to random selection, indicating no significant genomics support. Fig. 5e shows that RAMEN’s precision in detecting genomics-supported edges far exceeds that of random chance, as validated by the p-values from Binomial tests. Conversely, the edges missed by RAMEN showed no significant difference in support from the RNA data compared to randomly selected edges. These results affirm RAMEN’s effectiveness in uncovering relevant and supported edges overlooked by conventional statistical methods.

## Discussion

In this study, we address the challenge of mapping the complex network of relationships among numerous clinical variables and their effects on disease outcomes, such as COVID-19 severity and long COVID. Traditional correlation analysis methods often overlook many indirect relationships, and Bayesian network approaches, while insightful, are constrained by their high computational demand (Bayesian network structure learning is NP-hard). To overcome these limitations, we introduced RAMEN, an efficient method for Bayesian network structure learning aimed at uncovering complex interactions between clinical variables using cohort health records data. By employing random walks and a Genetic Algorithm, RAMEN effectively searches for the optimal Bayesian network structure, revealing the complex web of relationships among a wide range of variables. This method was applied to two distinct COVID-19 patient cohorts from the BQC19 dataset to delineate networks associated with COVID severity in hospitalized patients and long COVID in outpatient cohorts with repeated visits. RAMEN identified key variables linked to COVID severity and long COVID, corroborating with findings from existing research. The relationships identified by RAMEN were further validated through multi-omics data analysis within the BQC19 dataset, demonstrating the method’s capability to detect complex relationships and offering insights into disease mechanisms and outcomes.

RAMEN distinguishes itself from existing methods through its ability to efficiently identify directed relationships between variables, thereby clarifying causal pathways and differentiating variables as either drivers or effectors of disease outcomes. This method enriches the analysis by discerning both direct and indirect influences on disease outcomes, shedding light on the complex web of interactions that underlie disease mechanisms. Utilizing innovative random walks and Genetic Algorithms, RAMEN overcomes the computational limitations that beset conventional Bayesian network methods, demonstrating its ability to process large datasets, such as the BQC19 dataset with 880 variables, and deliver optimized network structures swiftly. This efficiency showcases RAMEN’s prowess in handling voluminous data effectively. Moreover, RAMEN introduces an approach in its network analysis by incorporating the disease outcome as a “terminal absorbing node.” This strategy makes all relationships within the network conditional on reaching the disease outcome, offering a nuanced perspective that prioritizes the significance of network paths leading to the disease outcome over mere direct correlations. This strategy is particularly effective in identifying variables that significantly influence disease outcomes, providing an analytical depth not available in most existing methodologies. Additionally, RAMEN pioneers in validating inferred Bayesian networks using multi-omics data, leveraging the growing availability of such data to delve into the biological mechanisms behind diseases. This approach marks a considerable leap in the systematic examination of reconstructed networks, effectively bridging the gap between computational analysis and biological validation. Through these advancements, RAMEN not only addresses the computational and analytical challenges posed by previous methods but also enhances the understanding of complex disease interactions.

RAMEN introduces a comprehensive toolset that fills gaps in current methodologies, enabling detailed analysis of complex relationships among clinical variables in diseases like COVID-19. This capability is pivotal for identifying biomarkers that facilitate early and accurate diagnosis, potentially leading to timely interventions and focused care by healthcare professionals to prevent complications. For instance, recognizing early joint pain as an indicator of increased risk for long COVID suggests that patients with such symptoms may benefit from prompt measures to mitigate the risk of prolonged illness. RAMEN’s strength lies in its efficient and precise exploration of the interactions between clinical variables and their impact on disease outcomes. This is expected to significantly improve the strategies for early diagnosis and intervention in severe and long-term COVID cases, thereby reducing the socioeconomic burden associated with COVID-19, especially its long-term effects. While the application of RAMEN has been demonstrated primarily within the context of COVID-19 datasets in this study, the framework is designed with the flexibility to examine the network of relationships among clinical variables across various diseases.

## Methods

### Datasets

In this study, we utilized multi-modal data encompassing clinical variables, RNA sequencing (RNA-seq), and SomaScan 5K analysis from two distinct cohorts of COVID-19 patients (hospitalized and outpatients) within the BQC19 dataset. 380 of these patients have undergone profiling for gene expression (via RNA-seq) and protein levels (using SomaScan[84]). The clinical data collected for these COVID-19 patients encompass a range of categories, including symptoms (such as cough and muscle pain), comorbidities (like asthma and diabetes), demographic information (sex and age), laboratory test results (including white blood cell counts and ALT levels), among others. For the purpose of this analysis, we focused on hospitalized patients to examine the correlation between COVID-19 severity and various clinical indicators. This subgroup was chosen due to the relative completeness of their clinical records, particularly laboratory results, which are often missing in outpatient records. Conversely, the outpatient cohort, characterized by a higher frequency of follow-up visits, provided a valuable dataset for the investigation of long COVID.

Our analysis only included clinical variables documented within the first month of patient care, as these data points hold significant relevance for the early diagnosis of COVID-19. Consequently, the outpatient dataset incorporated a total of 1440 patients and 84 clinical variables. In addition to outpatient data, we also analyzed data from 2,018 hospitalized patients, encompassing 297 clinical variables, to construct a network model identifying markers indicative of COVID-19 severity. Further, besides the BQC19 data, we also incorporated independent COVID-19 data from the Lawson Health Research Institute, which categorizes patients into three groups: outpatients, ward patients, and ICU patients. This dataset facilitated a broader analysis across different care settings, encompassing 27 variables across 66 patients. These patients, drawn from a tertiary care system in London, Ontario, Canada, were screened and enrolled for the study, offering insights into a wide array of clinical variables categorized into symptoms, comorbidity, demographics, laboratory test results, and other relevant factors.

### Bulk genomics data preprocessing

The preprocessing of the BQC19 RNA-sequencing data involved a standardized bulk RNA-seq workflow that commenced with a combined quality control and trimming step to ensure the generation of highquality sequencing data. Initially, the raw FastQ files were subjected to FastQC (version 0.11)[85] to assess the quality of the sequencing reads, identifying potential issues such as low-quality sequences, adapter contamination, and other anomalies that could affect downstream analysis. Immediately following this quality assessment, fastp (version 0.20)[86] was employed to trim and clean the FastQ files. This step not only removed adapters and low-quality bases identified during the FastQC analysis but also ensured that the remaining reads met a high standard of quality for accurate alignment. After the quality control and trimming processes, the cleaned reads were aligned to the human reference genome hg38 using HISAT2 (version 2.2.1)[87], which efficiently produced SAM files containing detailed information about each read’s alignment to the genome. These SAM files were then converted into BAM format using SAMtools (version 1.10)[88], optimizing the data for reduced storage space and faster access, which is crucial for efficient downstream processing. Following alignment, the featureCounts (version 2.0.1) [89] software quantified the aligned reads by counting the number of reads mapping to each genomic feature, such as genes or exons, generating a comprehensive raw counts file. This file forms the basis for analyzing gene expression levels across the samples. The final step in the data preprocessing involved normalizing and transforming the raw counts data using the DESeq2(version 1.38) [90] R package. This normalization is critical for adjusting for library size variations and sequencing depth differences across samples, ensuring that the gene expression data are comparable across the entire dataset. DESeq2’s normalization method prepares the data for accurate and meaningful downstream analyses, including differential expression studies, by providing a normalized set of gene expression values ready for in-depth biological interpretation. This integrated approach to RNA-seq data preprocessing—combining initial quality control with trimming, followed by alignment, quantification, and normalization—ensures that the BQC19 dataset is of the highest quality for subsequent analyses, laying the groundwork for insightful discoveries into the molecular mechanisms of COVID-19.

We also obtained SomaScan data corresponding to the circulating proteome measured by a multiplex SOMAmer affinity array (SomaLogic, 4,985 aptamers) from BQC19. SomaScan is a biotechnological protocol commercialized by the SomaLogic company (10). The SomaScan protocol comprises several levels of calibration and normalization to correct technical biases. Log2 and Z-score normalization were performed on each aptamer separately in addition to the manufacturer’s provided normalized data (hybridization control normalization, intraplate median signal normalization, and median signal normalization). Since the data was analyzed by SomaLogic in two separate batches, we applied the z-score transformation separately to each batch, to reduce batch effects. These additional transformations ensure that the measured values of different aptamers are comparable and can be used in cluster analysis.

### Clinical data preprocessing

In the preprocessing of the BQC19 clinical variable dataset to make it suitable for subsequent models, several steps were undertaken. Firstly, with a focus on early diagnosis, only the data from the first visit of patients with multiple visits were retained, although follow-up visits were considered for determining the presence of long COVID. For severity, for some patients, there has been more than one visit to the hospital in the first few days of showing symptoms, hence there are some measurements, such as “Temperature”, that were taken during both visits. Such measurements are kept in the dataset as “Temperature.1”. These variables can still be insightful, as they provide information during the early stages of symptoms of COVID-19. Secondly, variables exhibiting substantial missing values were removed. This decision was informed by creating histograms (as shown in Supplementary Fig. S14 and S15) to assess the distribution of patients having data for at least a certain number of variables, leading to the establishment of a threshold of 750 non-null values. Variables falling below this threshold for long COVID and COVID severity were excluded. Thirdly, for real-value clinical variables, discretization into bins was performed to simplify mutual information computation, with categorical variables being digitized (e.g., 0 for “no,” 1 for “yes”). This approach not only facilitates mutual information analysis but also ensures a standardized preprocessing for both clinical variables and RNA-seq and SomaScan data, as described in the standardized pipeline previously outlined.

### Mutual Information Calculation

To calculate mutual information effectively, it’s essential to ensure that the data pairs between the two variables’ vectors are both non-null. In our dataset, a common scenario is that a patient may have a recorded value for one variable but a null entry for another. Considering the independence of our patients’ medical histories, employing imputation to fill these gaps is not deemed rigorous for our study. Imputation could introduce inaccuracies into our dataset. To address the missing values issue in mutual information computation, we adopted a strategy of excluding any patient data with missing entries in either of the two variables under consideration. This was accomplished through numpy array manipulation techniques applied to the two vectors. In instances where the exclusion of missing data results in an absence of patient data for the two variables (a rarity), we default to assigning a mutual information value of 0. This approach allows us to calculate mutual information between two variables accurately, notwithstanding the presence of missing data.

### Build initial network structure with an absorbing random-walk-based permutation test

After the preprocessing stage, our methodology unfolds by creating a directed graph that encapsulates all clinical variables as nodes, with the connections—or edges—between these nodes weighted by the mutual information that quantifies the shared information between each pair of variables. Initially, this graph is fully connected, ensuring that each pair of nodes is linked, thus offering a comprehensive map for exploring potential relationships among all variables. To evolve this graph into a Bayesian network, which accurately reflects the true dependencies among clinical variables, we adopt a strategic approach that merges random walks with permutation testing to discern and eliminate less significant edges. This nuanced method begins with random walks across the network to evaluate the connectivity and importance of the paths between variables, with a special focus on paths leading to critical nodes such as those representing disease severity or the occurrence of long COVID.

In this refined approach, a random walk starts from a selected node X and persists until it either concludes at a designated absorbing terminal node (e.g., Severity or Long COVID) or surpasses a predetermined number of steps. A walk is categorized as “successful” if it ends at the absorbing terminal node, underscoring paths of potential significance in the disease’s mechanistic understanding. The transition probability from node X to another node Y during a walk is determined by the mutual information between X and Y, normalized to ensure the sum of probabilities to one across all potential next-step nodes from X. This probabilistic framework prioritizes transitions between nodes with higher mutual information, effectively highlighting stronger relationships.

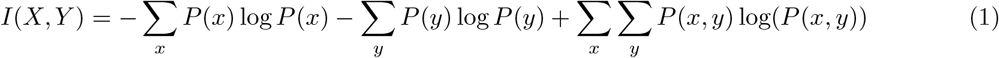

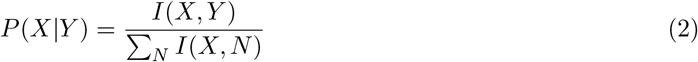

Concurrently, a permutation test is applied to assess the significance of these connections robustly. By randomly permuting the values among nodes and recalculating mutual information scores for these permuted networks, we generate a baseline distribution of mutual information under the hypothesis of no meaningful association between variables. Comparing the actual mutual information values against this distribution allows us to identify and subsequently trim edges that fail to statistically differentiate from what might be expected by chance. This combination of random walks for network exploration and permutation testing for evaluating edge significance presents a data-driven methodology to infer a Bayesian network. The result is a meticulously pruned network, conserving only those edges that are most indicative of genuine dependencies among clinical variables, thereby paving the way for more profound insights into the complex network of relationships that underpin disease pathology.

### Bayesian network inference using Genetic Algorithm

In our study, we utilize a Genetic Algorithm (GA), a score-based structure learning method, to optimize a directed Bayesian network that captures the intricate relationships among clinical variables. This optimization builds upon an initial network framework established through random walks, which helps in identifying a substantive starting point by trimming insignificant edges. The GA is particularly suited for this task due to the NP-hard nature of Bayesian network structure learning, facilitating an efficient search for a locally optimal solution. The process begins with the creation of ten directed acyclic graph (DAG) candidates, known as parent networks. These DAGs are constructed to reflect the complex interrelations suggested by the initial network, derived from the outcomes of random walks. The GA then iteratively applies a series of genetic operations on these networks until convergence is reached. Crossover involves the random selection of pairs of networks, between which edges are exchanged. This operation promotes the exploration of new network structures by merging aspects of two parents. Mutation introduces variations within a network by randomly performing one of three possible modifications: adding one or two consecutive edges, removing one or two consecutive edges, or flipping the direction of one or two consecutive edges. This step is critical for maintaining diversity among the network population and preventing premature convergence to local optima. Scoring of each network, whether an original candidate or one newly generated through crossover and mutation, is performed using an entropy-based scoring function designed for Bayesian networks, based on mutual information, as introduced by deCampos [91]. To extend this foundational framework, we have added a regularization term specifically to constrain network complexity. This modification to the scoring function achieves a more holistic evaluation by not only leveraging the predictive accuracy inherent in the entropy-based measure of mutual information but also incorporating a penalty for excessive complexity within the network structure. The revised scoring function is crafted to ensure that the optimization process for Bayesian network configurations effectively balances the trade-off between model accuracy and structural simplicity. The extended BN scoring function is articulated as follows:

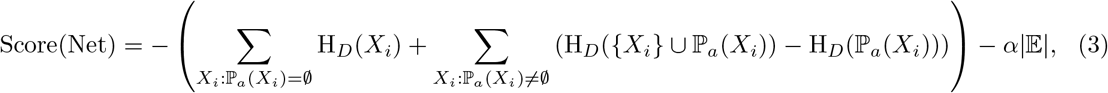

where *H*_*D*_ represents the entropy of a variable, reflecting its inherent uncertainty or informational content. 𝕡_*a*_(*X*_*i*_) denotes the set of parent variables for *X*_*i*_, and |𝔼| measures the network’s complexity via the total number of edges, thereby penalizing excessive complexity. The hyperparameter *α* modulates the impact of this regularization, striking a balance between the model’s simplicity and its fidelity to the data. This approach, inspired by the Kullback-Leibler (KL) divergence, strives to minimize the difference between the actual data distribution and the distribution implied by the Bayesian network model, ensuring the optimized network accurately reflects the data’s underlying structure. The RAMEN pseudo-program is as Algorithm 1.

### Computational validation of reconstructed network with the multi-omics data

In this section, we detail the computational approaches used to evaluate the networks formulated by RAMEN. **Validation through Clinical Variable Records:** To explore the associations between direct indicator variables and COVID-19 outcomes (severity of COVID-19 and long COVID), we constructed contingency tables that enumerate the patient counts for each pair of variable values. We applied Pearson’s Chi-square test to compute a p-value, thereby assessing the dependency between these variables. **Validation via Alternative Data Modalities (Gene Expression and Protein Levels):** We began by categorizing patients based on their variable values. We then identified Differential Expression (DE) genes among these patient groups using a t-test, as facilitated by the Python package diffxpy (FDR corrected *p <* 0.05 and | log_2_ fold change| *>* 0.6)[92]. To further confirm the correlation between the variables, we employed a hypergeometric test [93] to ascertain if there is a significant overlap in DE genes. In addition, to uncover potential biological functions linking the indicator to the outcome (either severity of COVID-19 or long COVID), we examined enriched pathways associated with these clinical variables by analyzing the list of DE genes using Toppgene. This analysis aims to deepen our understanding of the biological mechanisms at play.

### Support Vector Machine model to predict COVID Severity and long COVID outcomes

To evaluate the effectiveness of variables identified by RAMEN in predicting COVID outcomes, and to demonstrate the biological significance of these identified biomarkers, our approach involves a comparative analysis. Specifically, we aim to assess the predictive capability of variables selected by RAMEN against those identified through mutual information and correlation analysis. For this purpose, we utilized a Support Vector Machine (SVM) model with a linear kernel, implemented via the ‘svm’ module from ‘sklearn’[94], to predict disease outcomes based on three sets of variables: those identified by RAMEN, the top-ranked variables according to mutual information, and the top-ranked variables according to correlation. To ensure robustness and reliability in our findings, we conducted 20 separate experiments for each set of variables, yielding 20 data points per method. Each experiment comprised a 5-fold cross-validation procedure, facilitated by the ‘StratifiedKFold’ class from ‘sklearn.model selection’[95], to maintain the proportion of outcomes across folds. The performance of each model was evaluated based on the average prediction accuracy and F1 score across these experiments.

To statistically ascertain whether the predictive performance of RAMEN-derived variables was significantly superior to that of variables identified by mutual information and correlation analyses, we employed a one-sided Mann-Whitney U test, available through the ‘scipy.stats’ module. This methodological framework not only allows us to compare the effectiveness of RAMEN in identifying predictive biomarkers but also serves to validate the biological relevance of these biomarkers by their capacity to accurately forecast disease outcomes.

#### Algorithm 1 Refined Bayesian Network Optimization via Genetic Algorithm

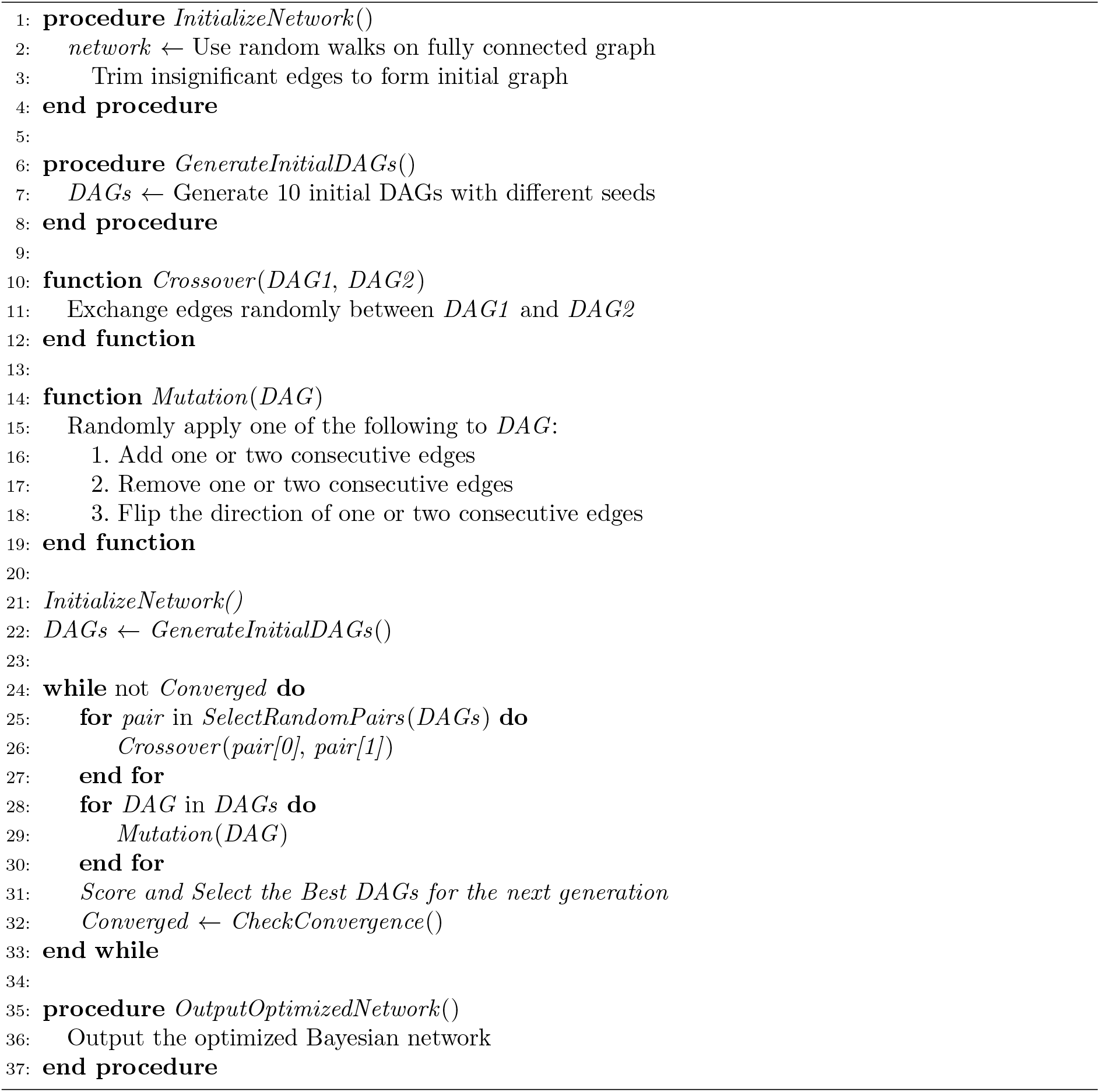

### Capturing feature importance via SHAP values

To determine the importance of variables identified by RAMEN in the SVM prediction model for COVID outcomes, we computed SHAP (SHapley Additive exPlanations) values, a technique based on game theory that assigns each feature a value indicating its contribution to the model’s prediction[96]. Following the training of the SVM model with RAMEN-selected features, we utilized the SHAP library to calculate the SHAP values for these features, thereby assessing their individual impact on model predictions. This computation was facilitated by an appropriate SHAP explainer, chosen to align with the SVM’s linear kernel. The resulting SHAP values were then aggregated to highlight global feature importance, providing a clear indication of how each variable influences the prediction outcome on average. By generating and analyzing SHAP summary plots, we could visually depict the rank and significance of each feature, thereby validating the predictive and biological relevance of the RAMEN-identified variables through their quantified contributions to the model’s predictive accuracy.

### Performance comparison via binomial test

To assess the significant differences between the proportions of supported edges identified by various methods, we employ a binomial test to calculate the p-value. Specifically, the test is formulated as *P* = 1 − binom.cdf(*k* − 1, *n, p*), where *n* represents the total number of edges analyzed, *p* is the proportion of supported edges observed in the method with the lower rate of support, and *k* is the actual number of supported edges corresponding to the higher proportion, expressed as an integer. Here, binom.cdf denotes the cumulative distribution function of the binomial distribution. A p-value less than 0.05 indicates that the method with the higher proportion of supported edges is significantly more effective at predicting edges that are corroborated by genomics data, thus affirming a statistically significant difference in performance between the two methods being compared. This statistical approach allows us to rigorously evaluate and demonstrate that RAMEN’s predictions are significantly more supported by genomic data compared to those derived from correlation analysis. p-values in Fig. 5 were calculated using this described approach.

## Software availability

The RAMEN software, including its source code and comprehensive documentation, is freely accessible online. Users can download and explore the software from the following URL: https://github.com/mcgilldinglab/RAMEN. For the purpose of visualizing networks reconstructed by RAMEN, Cytoscape[97], a prominent tool for network visualization, was employed. Additionally, an interactive web portal has been developed to enhance the accessibility and usability of the reconstructed networks. This portal allows users to dynamically interact with the networks generated by RAMEN and can be accessed at http://dinglab.rimuhc.ca/pgm/.

## Acknowledgement

This work was partially supported by CIHR PJT-180505, FRQS 295298, 295299, and NSERC RGPIN2022-04399 to JD. We thank the BQC19 initiative for granting us access to the multi-omics COVID data.

## Notes

http://dinglab.rimuhc.ca/pgm/

https://github.com/mcgilldinglab/RAMEN

### Competing Interest Statement

The authors have declared no competing interest.

### Summary of Updates

The new version features additional benchmarking which better showcases the rigorousness of the method. Additionally, we have provided more figures that will give the reader a better understanding of the RAMEN method. There is also a new independent study with the RAMEN method on an additional dataset provided by the researchers at the Lawson Health Research Institute.

