## Supplementary Figures for "RAMEN Unveils Clinical Variable Networks for COVID-19 Severity and Long COVID Using Absorbing Random Walks and Genetic Algorithms"

### 1 Supplementary Figures

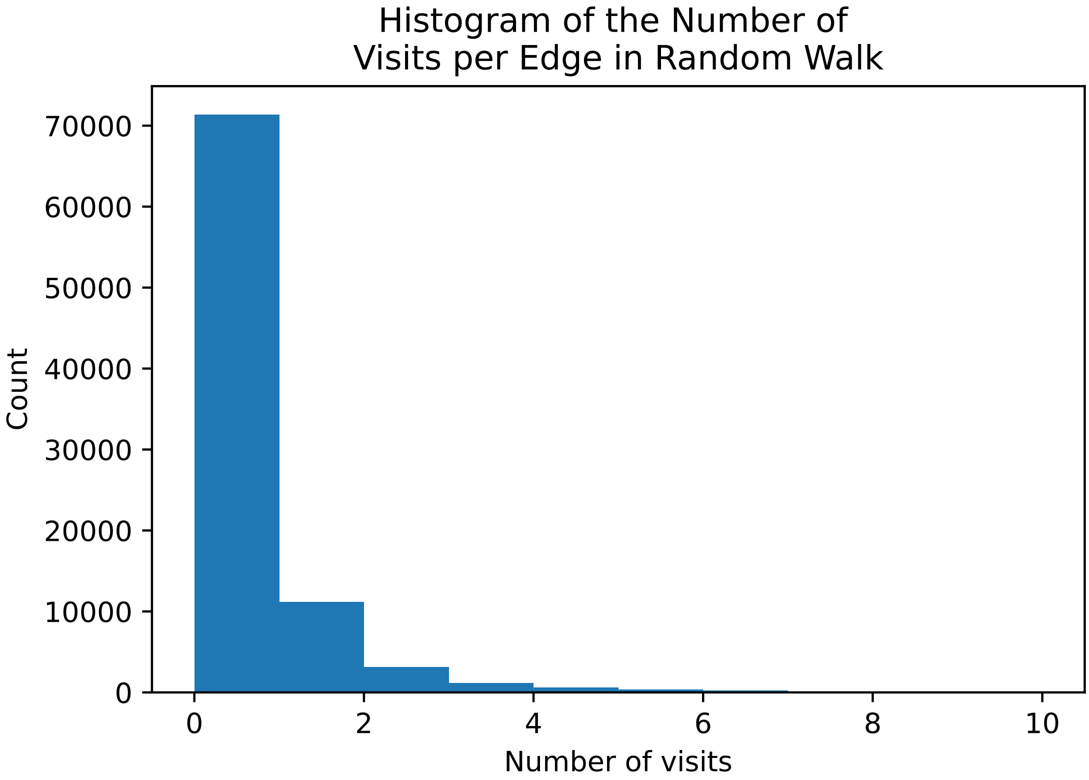


**Fig. S1**: **This figure represents the histogram of the number of visits per edge in the random walk where the edge weights are shuffled to create the background distribution.**

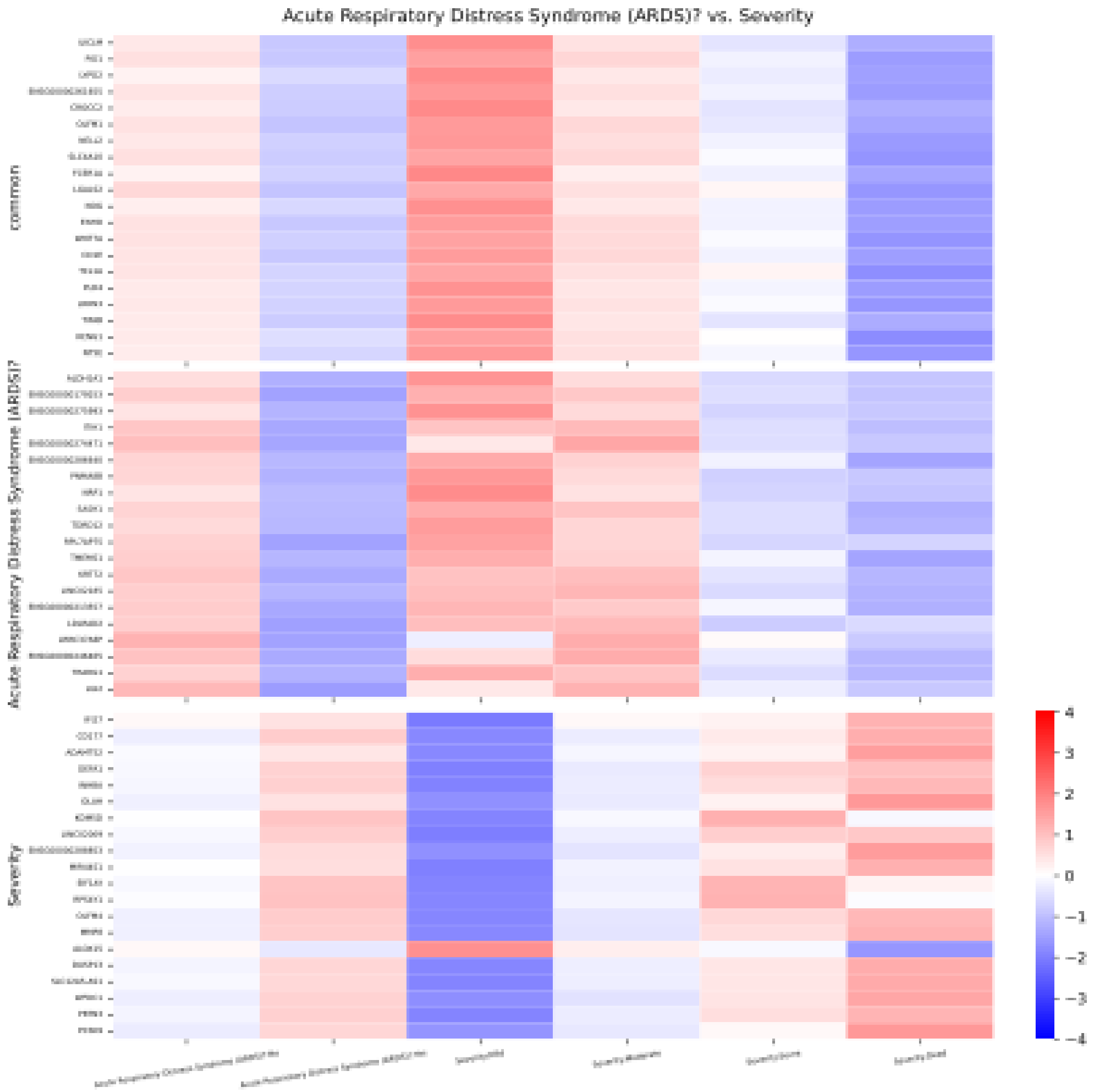


**Fig. S2**: **Examining the edge connecting Acute Respiratory Distress Syndrome (ARDS) and severity in the reconstructed COVID severity network with RNA sequencing data.** All differential genes associated with ARDS (i.e., genes whose expression changes among the patients with different levels of ARDS and the differential genes associated with COVID severity are identified. Then we visualize the following three groups of genes: (1) 'Common' shows the shared genes between those two groups of genes; (2) 'ARDS' shows the unique differential genes for the ARDS variable; (3) 'Severity' shows the unique differential genes for COVID severity.

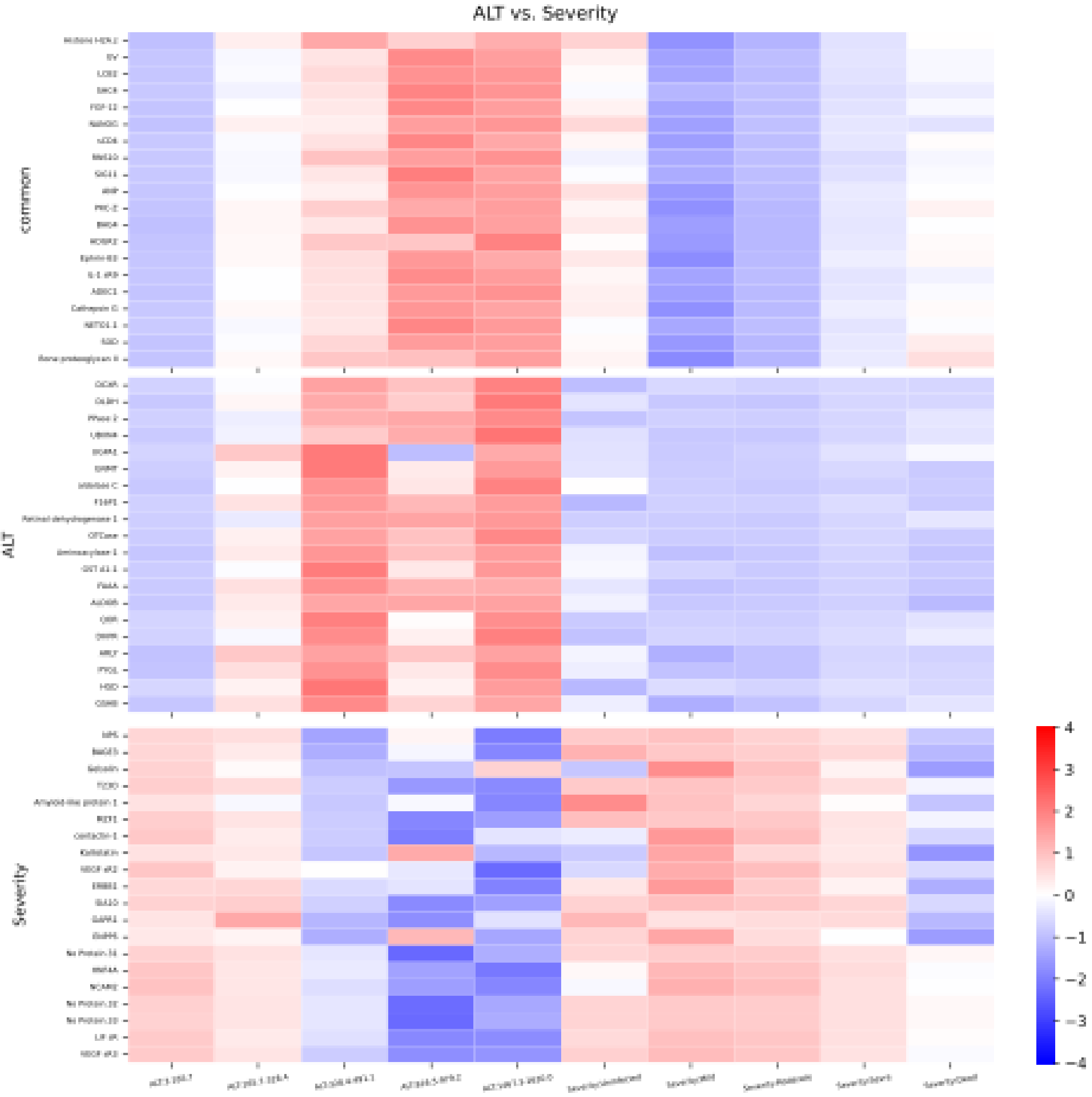


**Fig. S3**: **Examining the edge connecting Alanine Transaminase (ALT) and severity in the reconstructed COVID severity network using Somascan (proteomic) data.** The heatmap shows three groups of proteins: (1) The common proteins between the proteins associated with ALT and Severity (2) The proteins that are associated with ALT, but no Severity (3) The proteins that are associated with Severity, but not ALT.

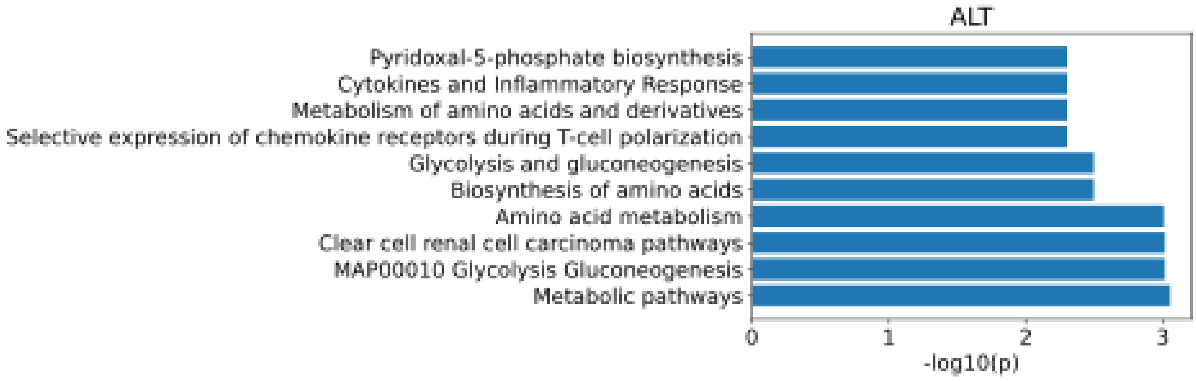


**Fig. S4**: **Enrichment Analysis for ALT using Somascan (proteomic) data for the 10 top pathways.**

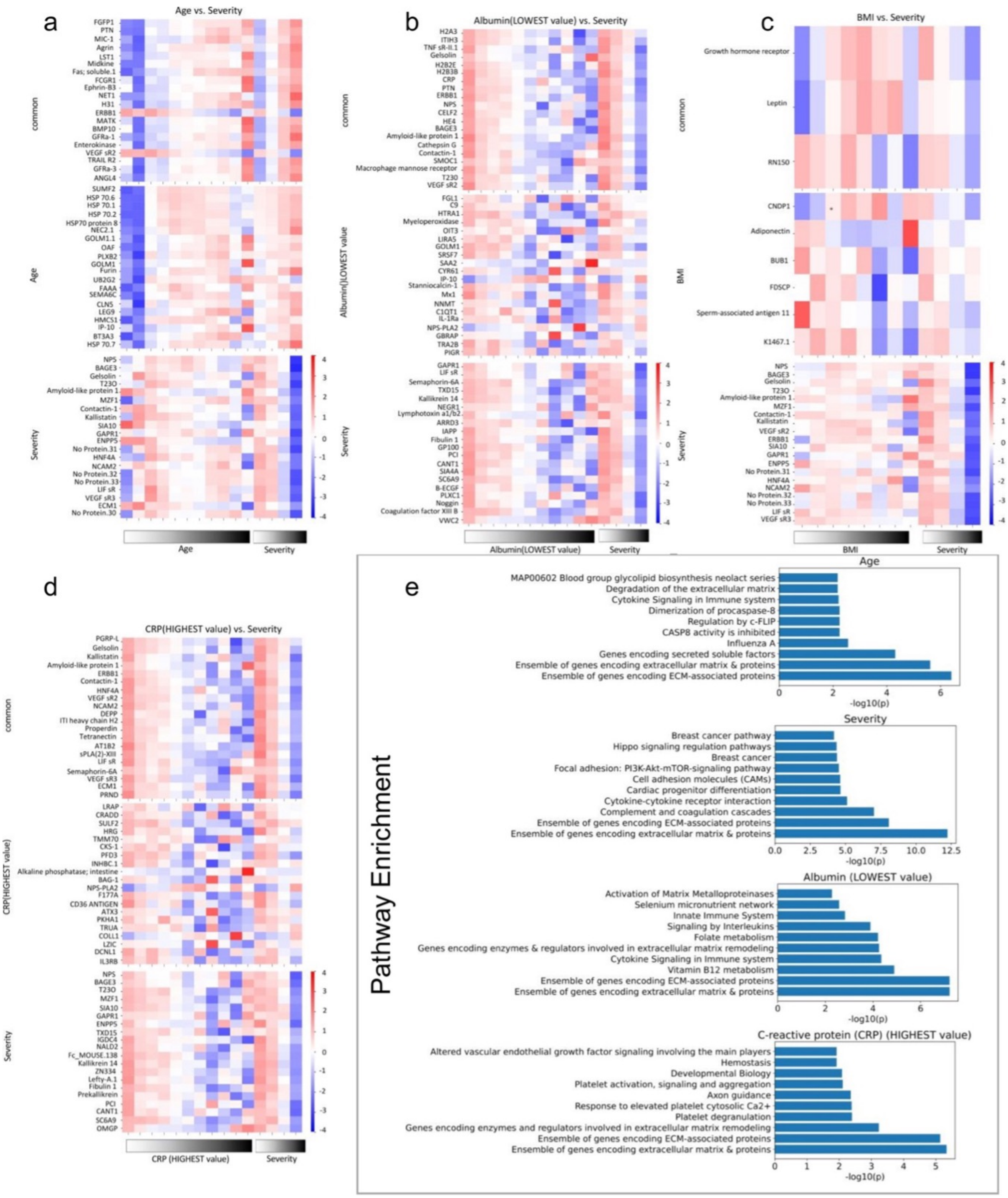


**Fig. S5**: **Somascan (proteomic) performing DE analysis for the variables connected to severity in the COVID-19 severity network**. The meanings are the same as supplementary Sup. **??** and Sup. We generated heatmaps for top 100 protein coding differentially expressed genes (ranked by fold changes) for four important variables related to Severity and showed their overlap with Severity. We also performed pathway analysis and showed the top 10 pathways for the 4 studied variables. **??**. **a** Age **b** Albumin **c** BMI **d** CRP **e** Enrichment analysis from the genes coding top 100 DE protein (ranked by fold changes) for the variables in **a**, **b**, **c**, **d**.

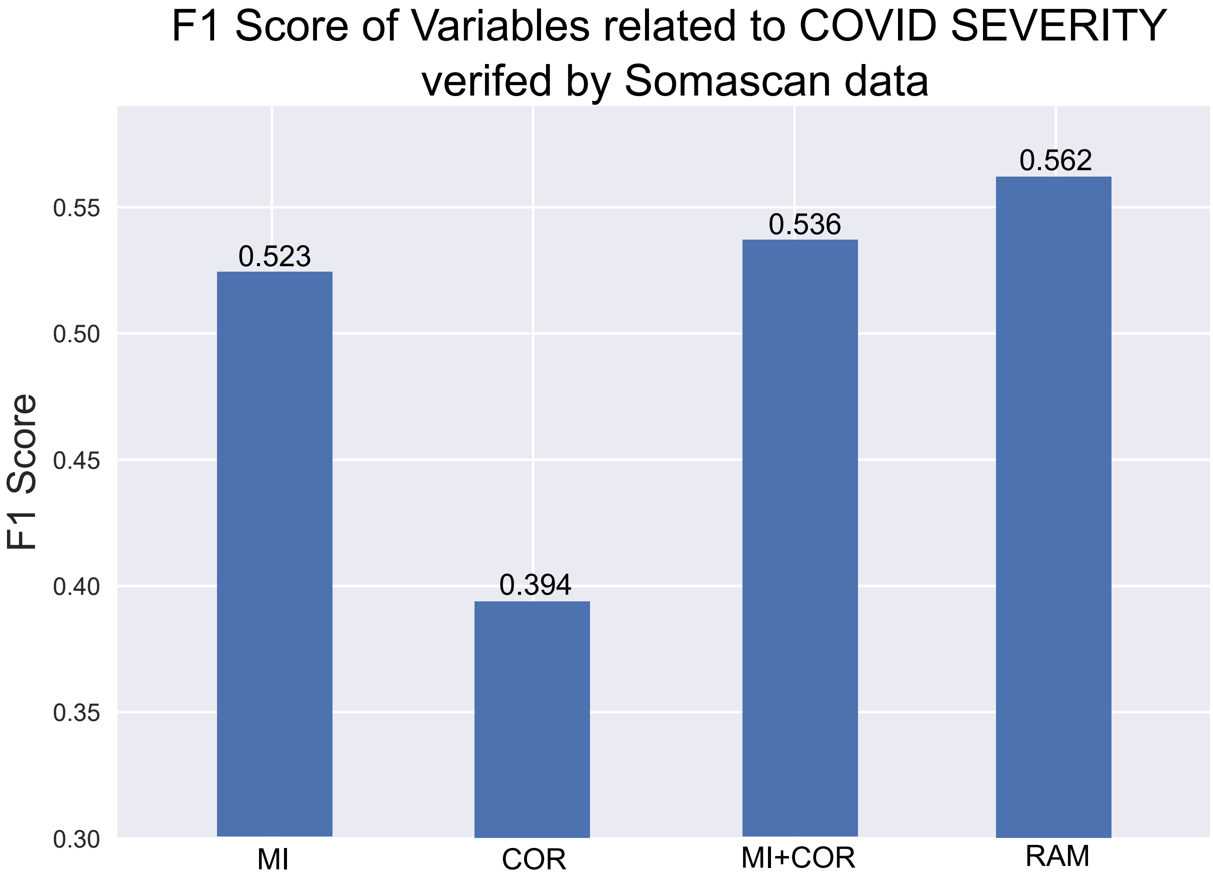


**Fig. S6**: Validation of COVID-19 severity indicators using various methodologies. Each method on the x-axis (MI: mutual information; RAM: RAMEN; COR: Pearson correlation) classifies variables into indicators or non-indicators, with Somascan data providing the basis for ground truth. A variable is considered an indicator if its DE genes significantly overlap with those associated with COVID-19 severity, assessed via a hypergeometric test. The classification performance of each method is quantified using the F1 Score from verifying the variables found by each method against the ground truth.

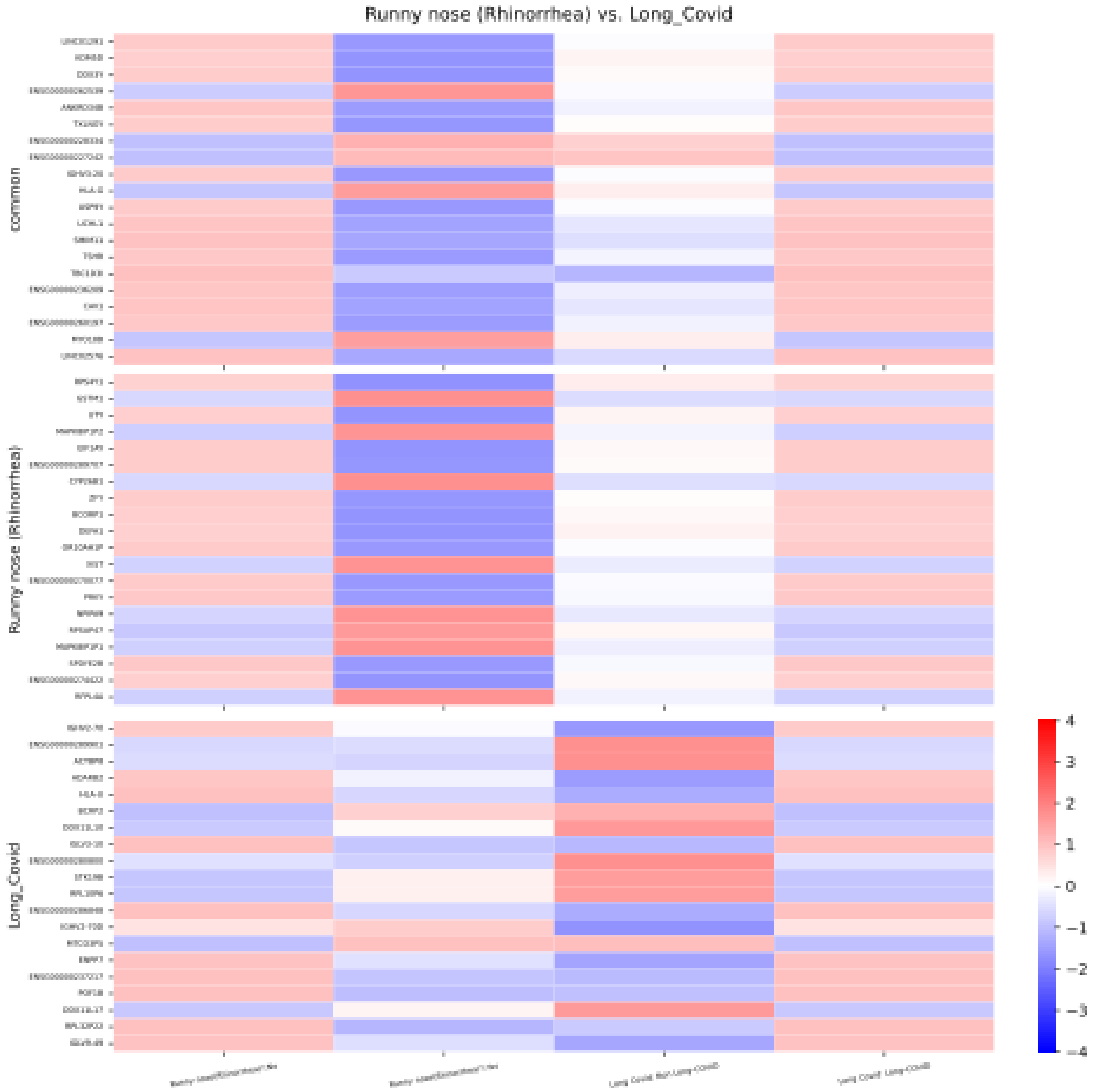


**Fig. S7**: **Examining the edge connecting Runny Nose and long COVID in the reconstructed long COVID network using RNA-seq data.** The heatmap is plotted in the same way as Supplementary Fig. S2.

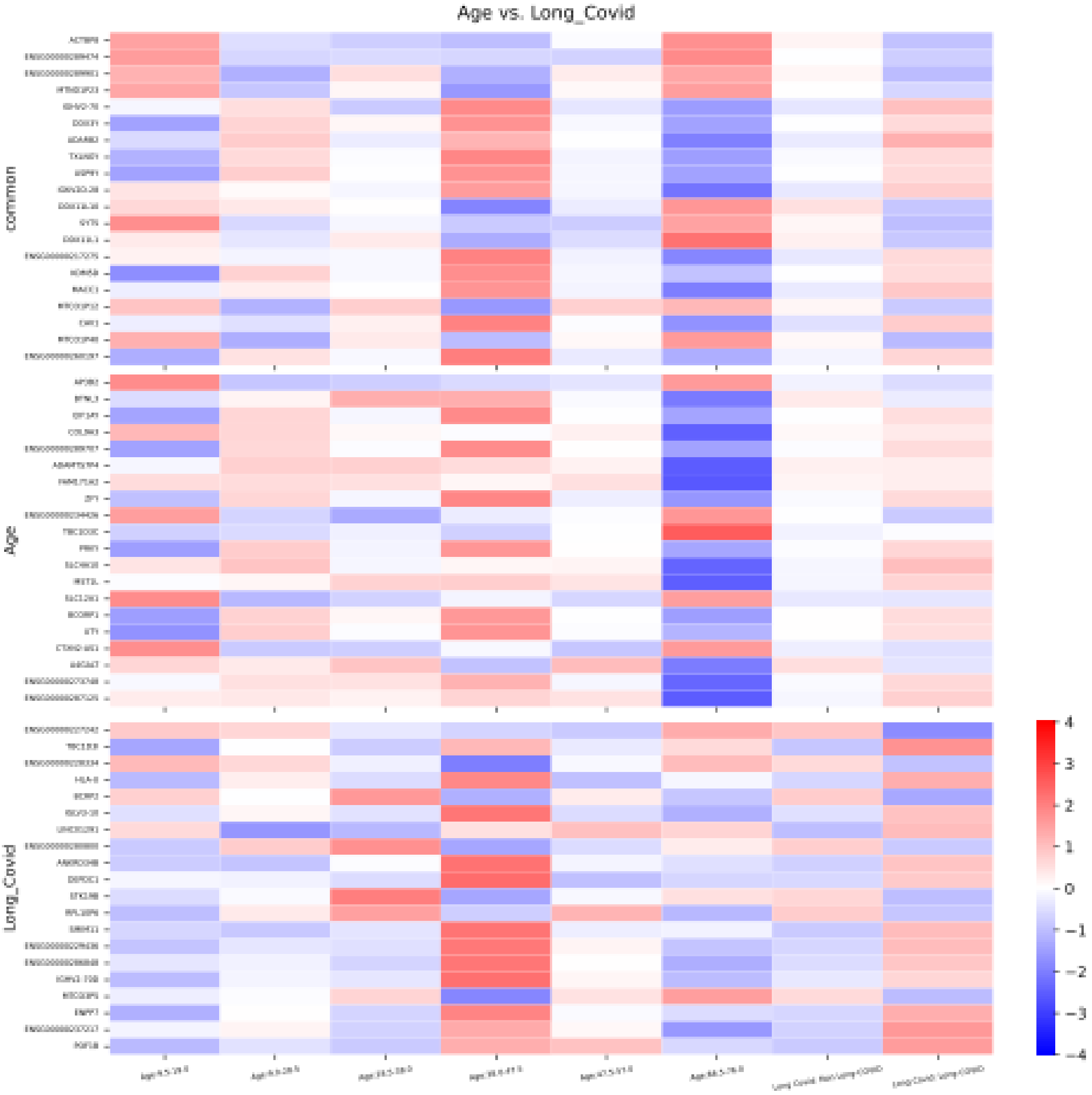


**Fig. S8**: **Examining the edge connecting Age and long COVID in the reconstructed long COVID network using RNA-seq data.** The heatmap is plotted in the same way as Supplementary Fig. S2.

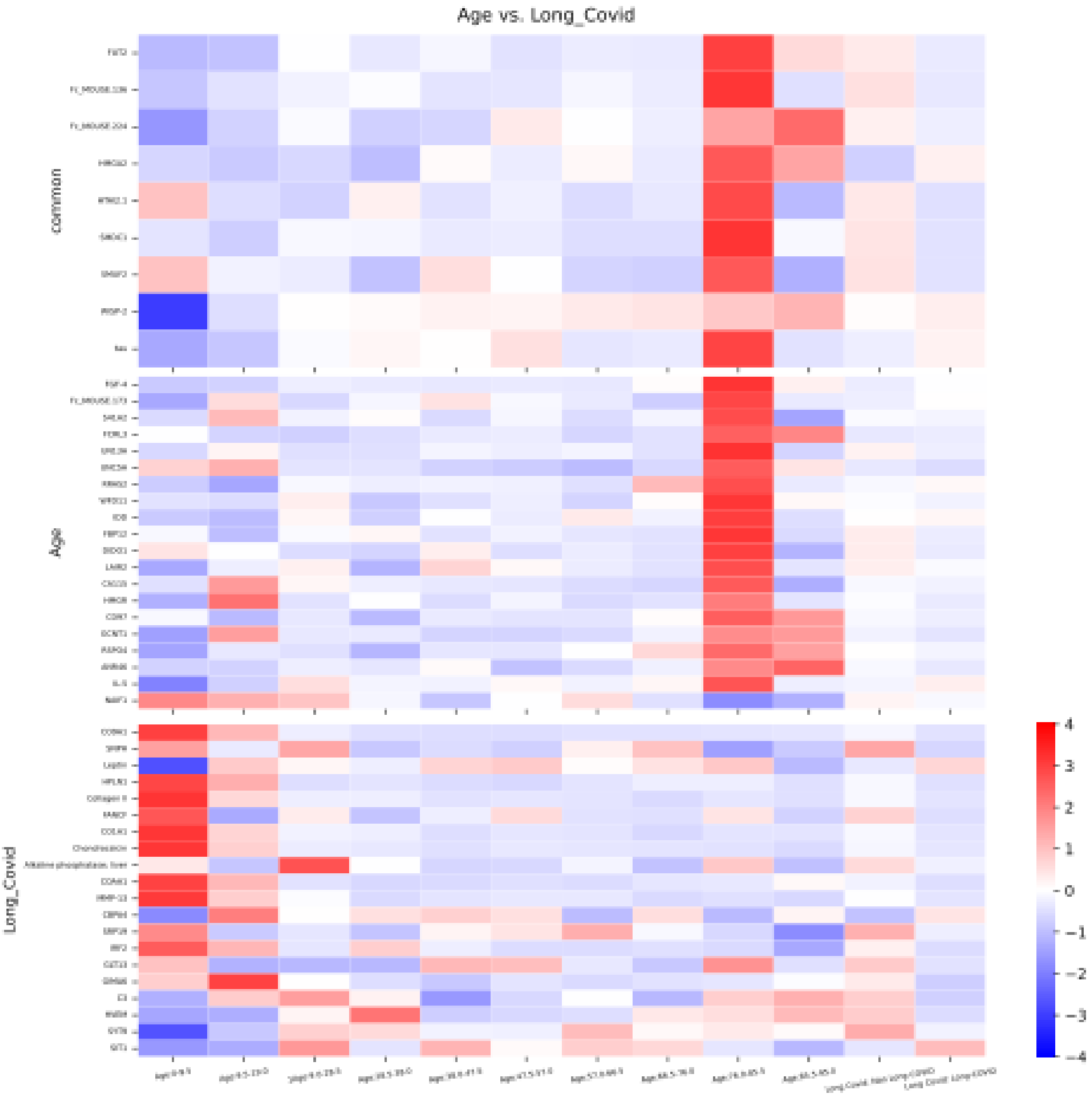


**Fig. S9**: **Examining the edge connecting Age and long COVID in the reconstructed long COVID network using Somascan (proteomic) data.** This heatmap is plotted similarly to the method described in Supplementary Fig. S2.

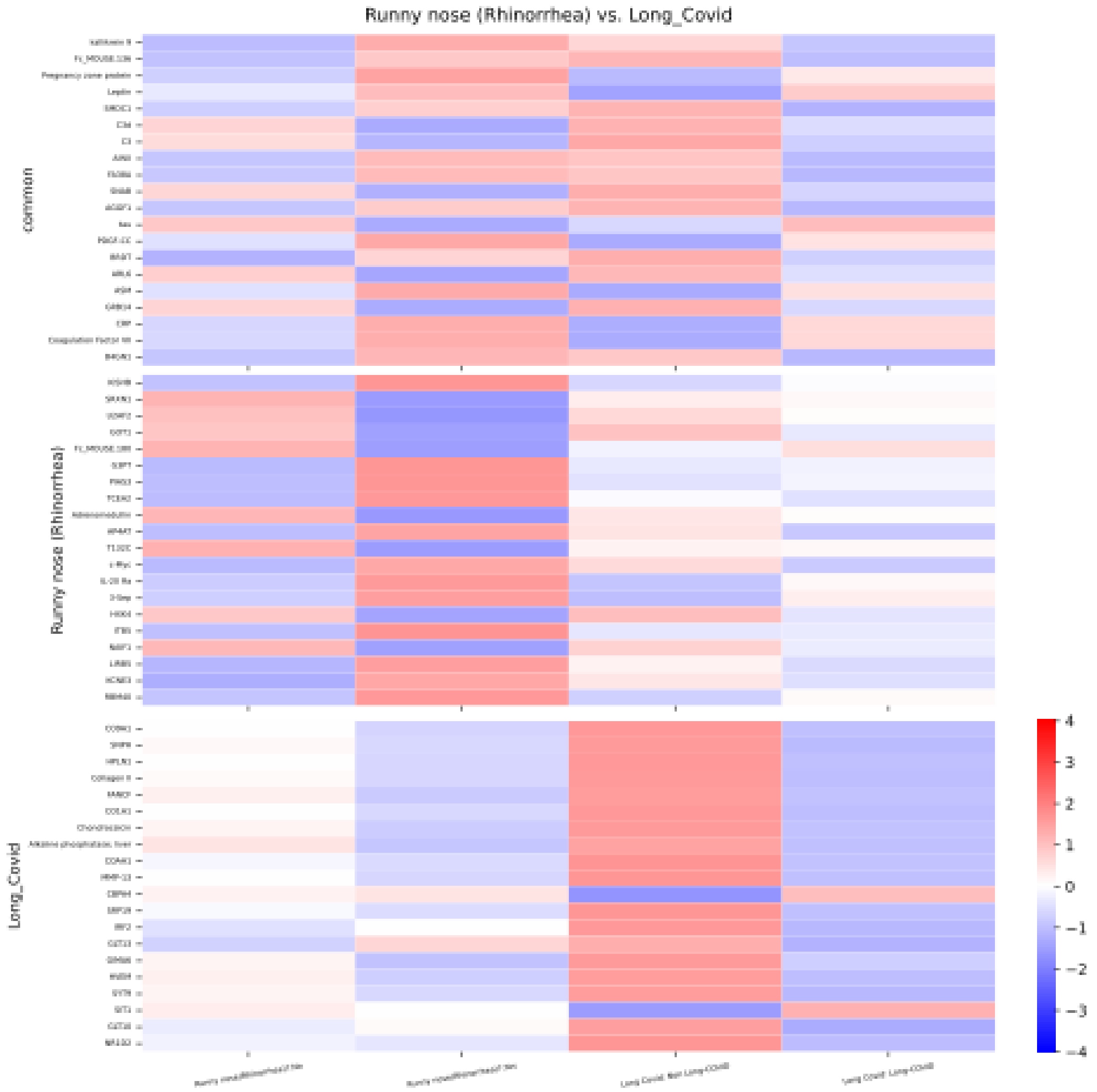


**Fig. S10**: **Examining the edge connecting Runny nose (Rhinorrhea) and long COVID in the reconstructed long COVID network using Somascan (proteomic) data.** The heatmap is plotted similarly to the method described in Supplementary Fig. S2.

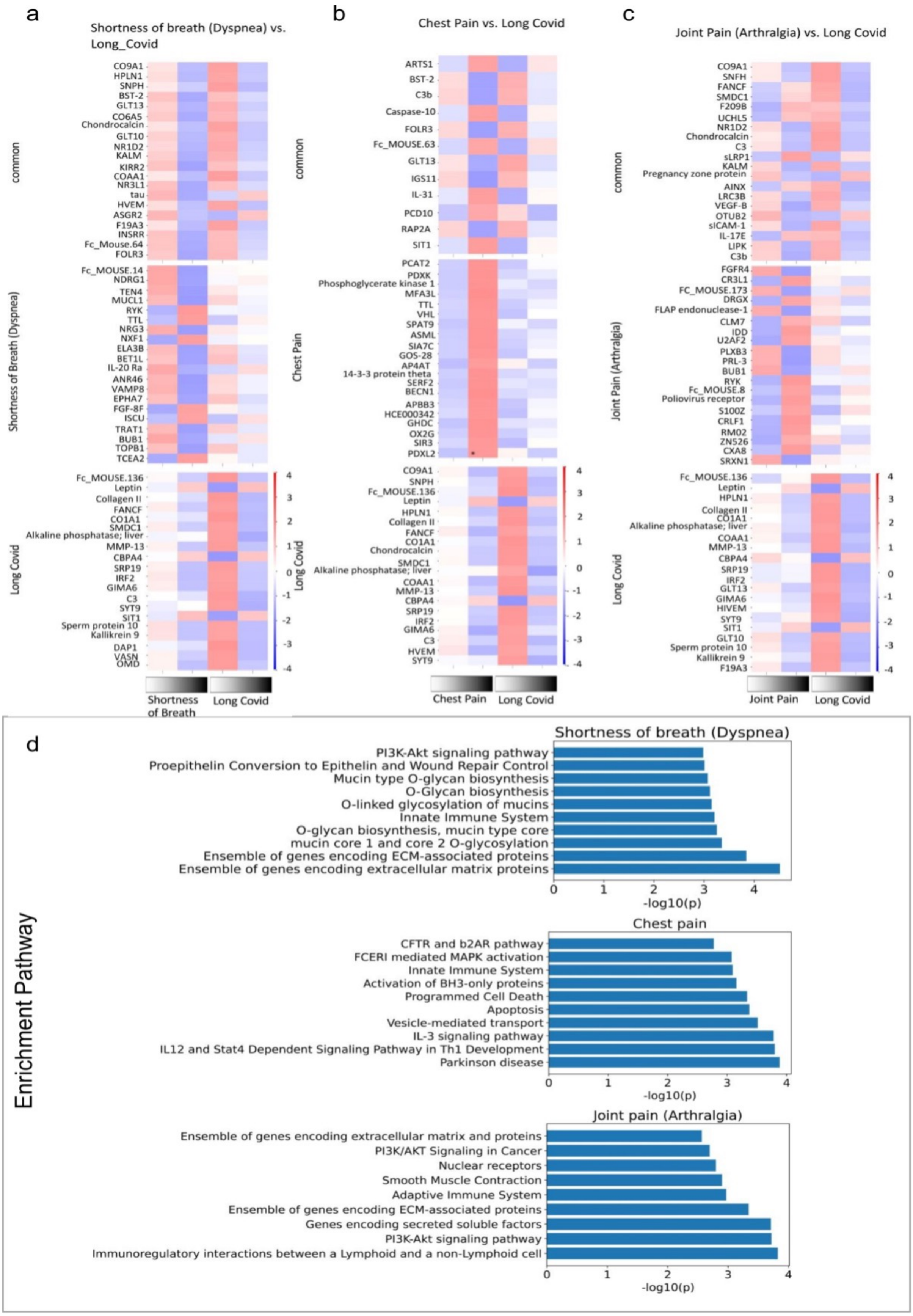


**Fig. S11**: **Validating network connections using multi-omics data.** The heatmaps show the protein expression pattern of pairs of variables connected/not connected in our long COVID network. In each plot, top genes are grouped into common top genes. We have also shown the top pathway enrichments for the 3 studied variables. **a** Shortness of breath (Dyspnea). **b** Chest pain. **c** Joint pain (Arthralgia). **d** Pathway enrichment for the previous variables.

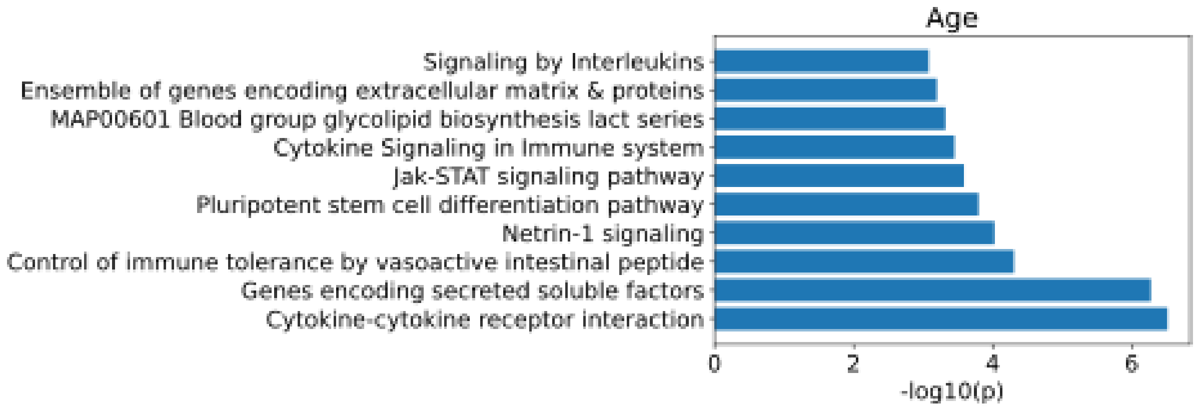


(a) Age

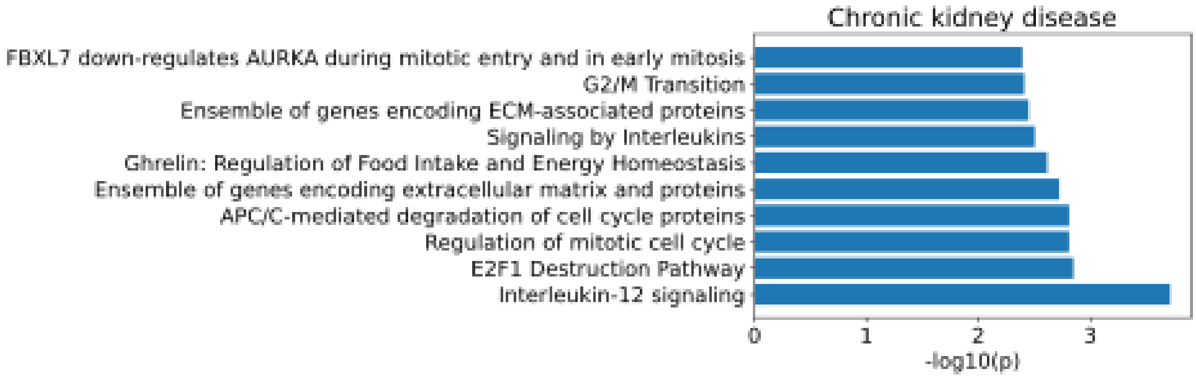


(b) Chronic kidney disease

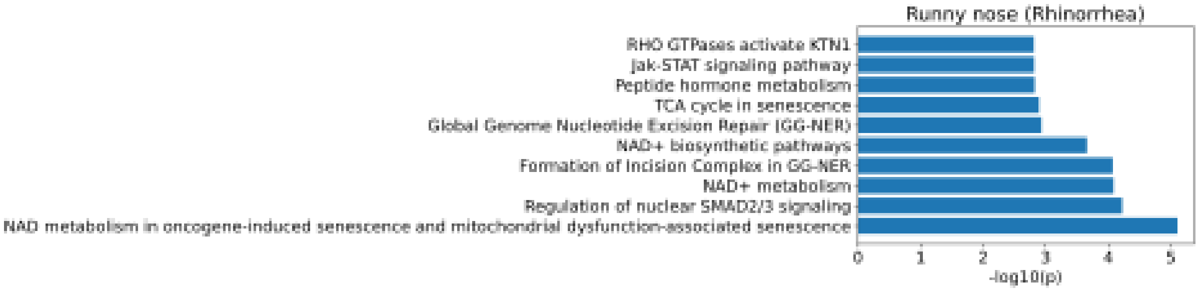


(c) Rhinorrhea

**Fig. S12**: **Enrichment Analysis using Somascan (proteomic) data for Age, Chronic kidney disease, and Runny nose (Rhinorrhea).** We have shown respectively their top 10 pathways.

| **Variable** | **p-value (chi-squared test)** |
| --- | --- |
| Shortness of breath (Dyspnea) | $2.946\,614\,013\,245\,262 \times 10^{-30}$ |
| Joint pain (Arthralgia) | $1.956\,442\,168\,076\,805\,4 \times 10^{-22}$ |
| Loss of taste / Lost of smell | $3.692\,012\,678\,509\,67 \times 10^{-18}$ |
| Chest pain | $7.894\,555\,024\,236\,59 \times 10^{-18}$ |
| Fatigue | $2.769\,582\,329\,243\,375\,4 \times 10^{-11}$ |
| Diarrhea | $6.557\,511\,286\,198\,923 \times 10^{-12}$ |
| Extremity weakness or numbness | $8.388\,045\,023\,351\,143 \times 10^{-11}$ |
| Number of comorbidities | $5.506\,442\,515\,645\,947 \times 10^{-6}$ |
| Nausea / vomiting | $5.805\,266\,435\,467\,504 \times 10^{-9}$ |

**Table S2**: **Chi-squared test results for various symptoms**

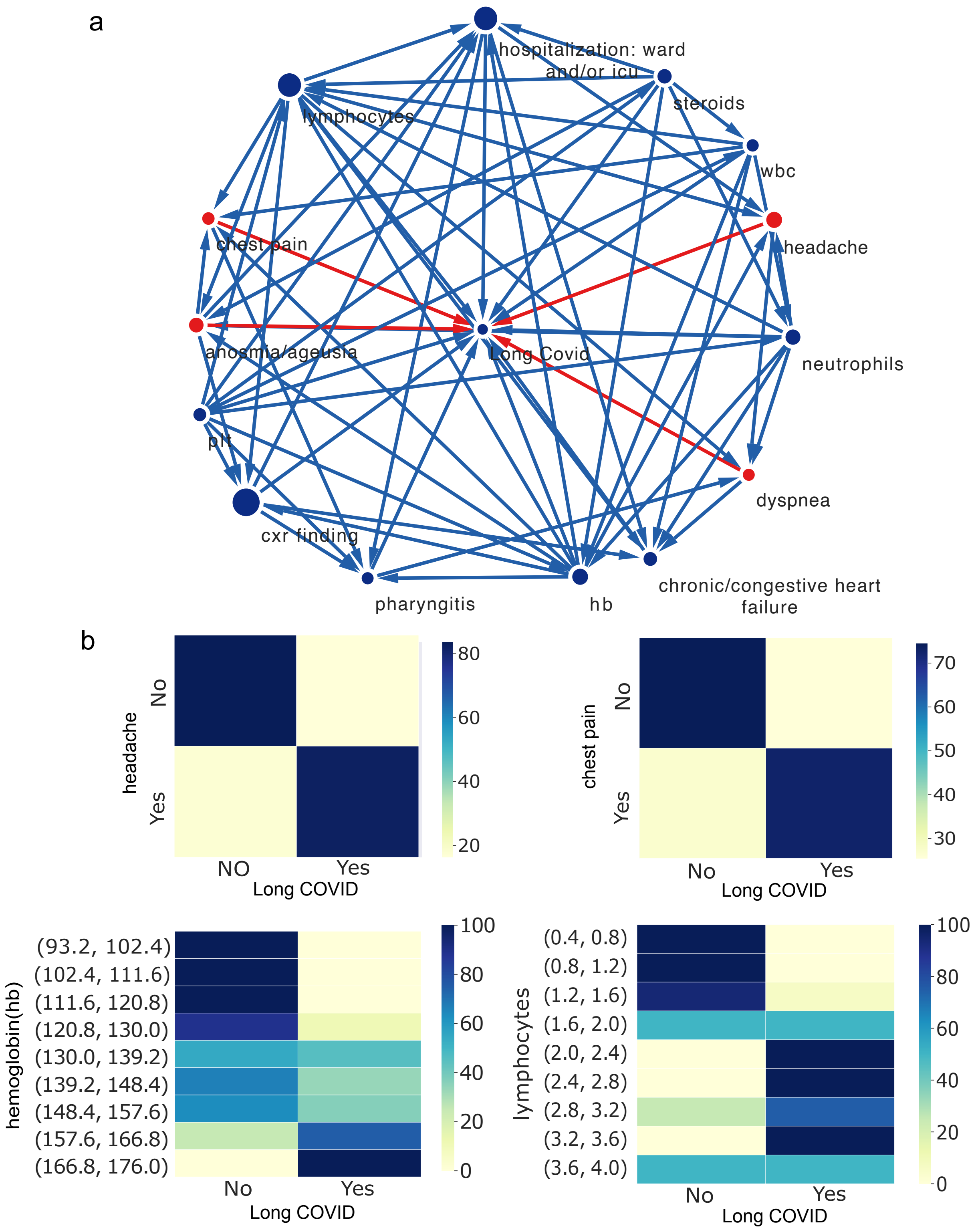


**Fig. S13**: **Constructing long COVID network using an independent dataset from Lawson Health Research Institute a,** The long COVID network constructed from the Lawson Health Research Institute's independent dataset, which includes 8 variables also present in the BQC19 outpatient data. Significantly, this study corroborates the identification of the same 4 direct long COVID indicators (emphasized in red) as found in the BQC19 analysis. **b,** Heatmaps showing the conditional distribution of the long COVID variable, conditioned on the values of the indicators found by RAMEN.

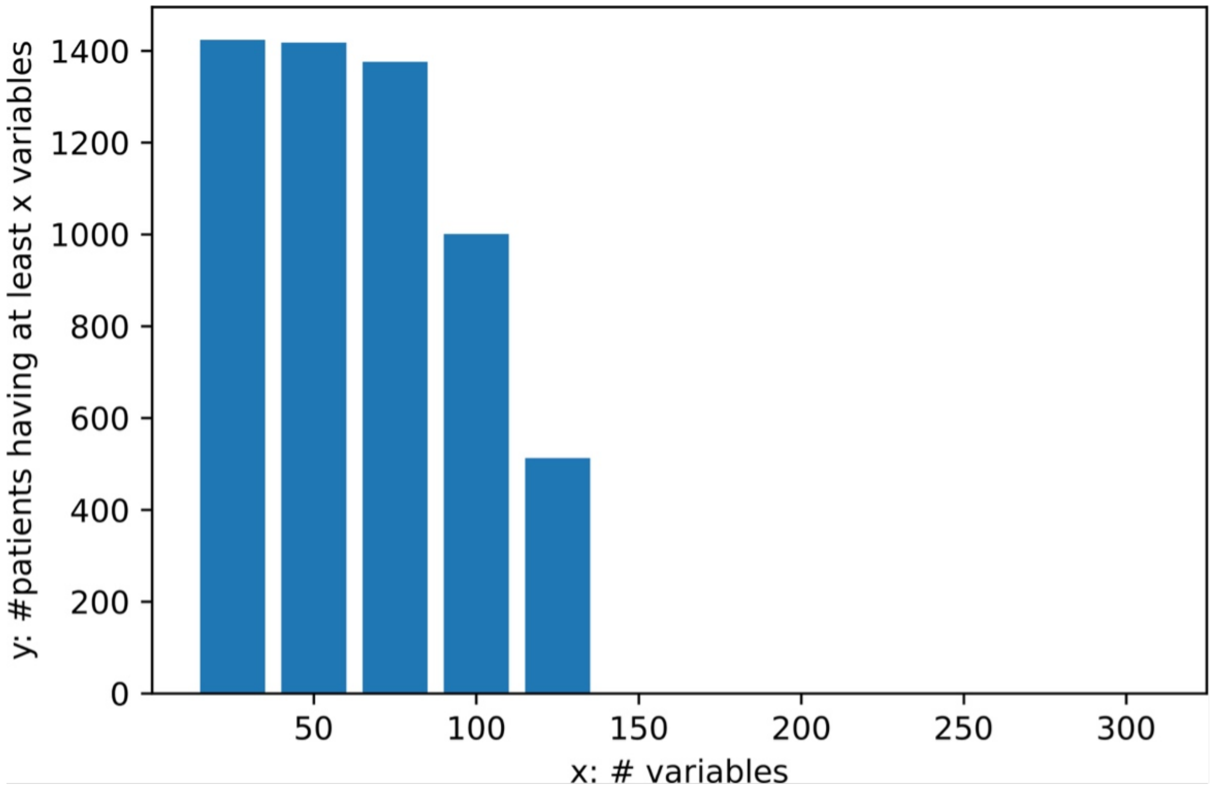


**Fig. S14**: **Number of variables having at least y number of recorded values in the out-patient dataset for studying long COVID.** We discard variables with fewer than 750 recorded values.

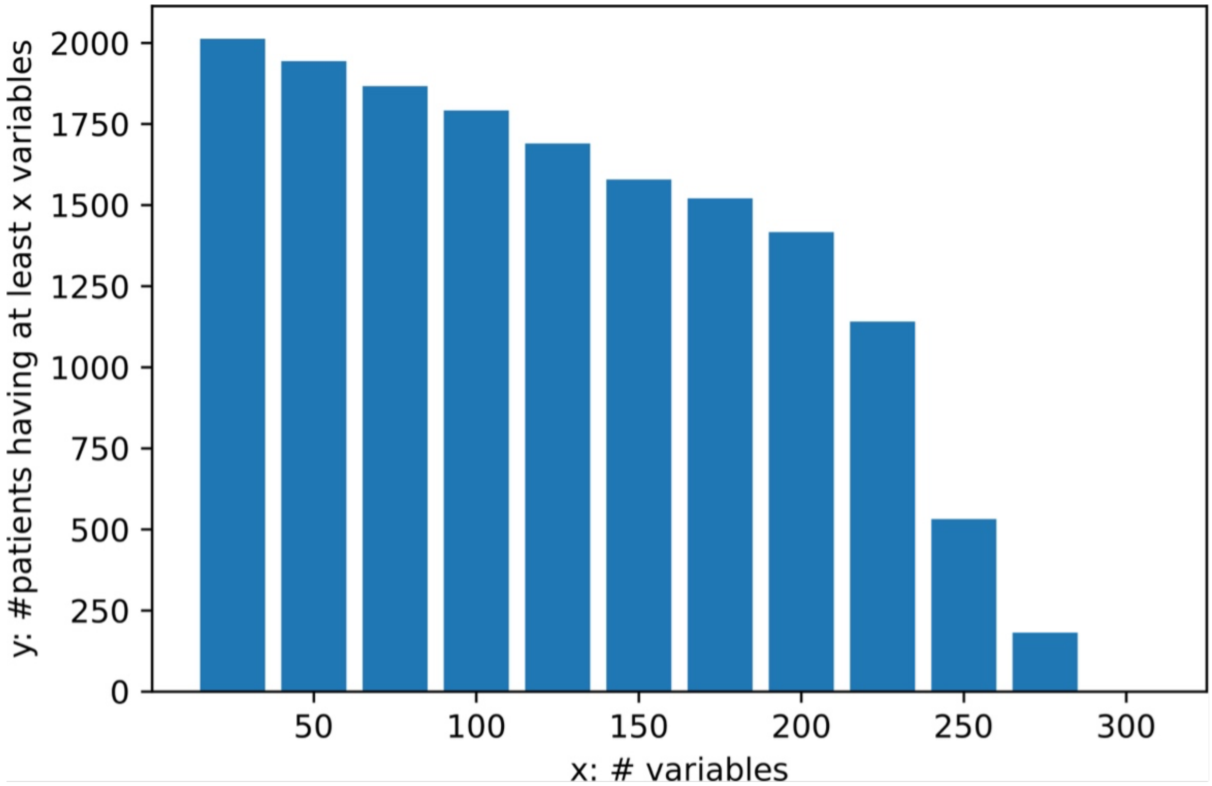


**Fig. S15**: **Number of variables having at least y number of recorded values in the hospitalized patients dataset for studying COVID-19 severity.** We discard variables with fewer than 750 recorded values.

### 2 Supplementary Methods

#### 2.1 Details about Data Processing

We create one data sample for each patient using the first visit. The subsequent visits of the patient are used to complete missing information for the given patient. If the value for a clinical variable is missing for a patient, information from the subsequent visits is checked to fill the missing values. The first visit contains the most information, as variables such as Weight, Height, BMI, and more are generally not taken for subsequent visits. If the value missing from the first visit cannot be found in subsequent visit as well, then it will remain a missing value for the analysis. As the study is about Severity and long COVID, all the negative patients are also removed. Following the initial re-formatting of the data, need to take care of the missing values of the clinical variables, which are very common in the BQC19 dataset and will unavoidably hampers the inference of the network. We solved this problem using two complementary strategies. First, clinical variables dominated with missing values (¡ 750 recorded values) will be removed from downstream analysis. The cut-off 750 is picked according to Fig 12 and 13. We pick an "elbow" value such that we keep a good number of variables and further including variable will sacrifice variable quality to a much larger extent. Second, the mutual information between two clinical variables is calculated only based on the patients with both variables. Furthermore, while a significant proportion of the clinical variables are binary or categorical, many other variables are real values (e.g., age or ALT levels), which significantly increase the complexity for the inference of their relationships. Therefore, to simplify the network inference, here we discretize the real value variables. For examples, patients across different age groups are also binned into 10 groups.

### 3 Supplementary Results

#### 3.1 COVID-19 severity network is supported by the COVID-19 hospitalized cohort

Heatmaps in Fig.3 show the percentage of patients with different severities for different value range of clinical variables. Namely, Fig.3 heatmaps indicates whether one clinical variable could suggest a special preference of COVID-19 severities. Here, we showed the heatmap for 5 top variables connected to COVID-19 severity-relevant variables (ALT, CRP -Highest value, Age, and Albumin, BMI). To demonstrate the specificity of RAMEN, we also show the heatmap for a clinical variable, not linked to COVID-19 severity-irrelevant variable (Leg swelling (Edema)). In general, for the positive variables that directly linked to COVID-19 severity in our reconstructed network, the distribution of the COVID-19 severity (% of patients across different severities) changes substantially accordingly to the value change of the positive variables. On the other hand, the distribution of the COVID-19 severity stays unchanged in response to the change of the negative variable. Fig.3 panel a clearly shows that the ALT is associated with COVID severity. An elevated level of ALT indicates increasing COVID severity. In fact, from an ALT value of 328.4 to 1979.2, 100% of patients have more than a moderate Severity. In the highest interval, 50% of patients have a mild reaction, while 50% of patients have a severe one. This could be explained by the rarity of patients that have an ALT value that is as high. Hence there might be some deviations from the general trend. From the heatmap shown in Fig.3 panel b, we can notice an increasing proportion of severe COVID patients along with the increase of C-reactive protein (CRP) (Highest value). Indeed, for patients with CRP between 0 to 63.8, only 12.2% of patients have a severe or worse reaction to COVID. Among patients with a CRP between 63.8 to 127.6, 23.51% of patients have a severe or worse COVID outcome. The heatmap clearly indicates a strong correlation between CRP (highest value) and COVID severity. The Fig.3 panel c suggest that the age is another important risk factor for severe COVID outcomes. From the heatmap, we can clearly see an increasing risk of more severe COVID outcomes along with the aging process. Specifically, the majority of young patients (patients below 21 years old) only show mild symptoms. However, more patients age between 21 and 31.5 develop more severe outcomes. The proportion of moderate, severe, and dead COVID infectants have all increased by a significant margin, 10%, 8% and 1.3% of patients respectively. Most patients aged 31.5 and above (¿75%) develop moderate or above COVID severity, which suggest most adult COVID patients develop at least moderate symptoms. The percentage of severe COVID patients (severe or dead) really ramps in among patients aged above 52.5. There are generally over 30% of patients aged between 52.5 and 84.0 developed severe or dead COVID outcome. It is worthy mentioned that we don't have sufficient patients aged 84.0 and above in our cohort and thus the stats associated with this age group is relatively limited.

The Albumin heatmap in Fig3. Panel d shows an inverse trend between the value of Albumin (Lowest value) and Severity. Indeed, as the value for minimum value of Albumin increases, the level of severity decreases. Starting from a minimum Albumin value of 15.2, the proportion of patients that are dead from COVID or have severe reactions is strictly decreasing, both eventually to a minimum of 0. At the same time, the proportion of patients that have moderate reactions increased from 0% at 10.1 up to a maximum of 73% at 35.6. Then it starts decreasing, but that is due to the increase to the proportion of mild reactions from an Albumin level of 25.4 to 56, eventually reaching 100%. These observations confirm that evaluated Albumin level generally indicates a less severe COVID outcome. The BMI heatmap in Fig.3 e demonstrated that a high BMI is another risk factor for severe COVID. In general, COVID infectants with higher BMIs is more likely to develop severe or worse (dead) COVID. Indeed, aside from the interval value of 60.8 to 66.9, the value of Severe reactions is strictly increasing with the increase of BMI. From the interval 42.5 to 48.6, the proportion of moderate is no longer dominant, and the proportion of severe reactions has surpassed and become the dominant reaction in the distribution. The death proportion also shows an increasing trend from BMI values of 12 to 48.6. Leg Swelling (Edema) is a negative variable that is not connected to severity in the reconstructed network by Ramen and serves as a negative example for comparison to the positive variables. From the heatmap shown in Fig.3, we clearly observe that the distribution of COVID severity stay almost identical among patients with and without Leg swelling (Edema). This suggests that Edema is very unlikely to be an important indicator for COVID severity and our model successfully captured this fact. long COVID network is supported by the BQC19 outpatient cohort Chest Pain and Join Pain are known symptoms related to long COVID, and our model has successfully captured that (Fig.6). Indeed, in the heatmap of both cases, we clearly observe that the proportion of patients with those two symptoms tend to develop long COVID. Specifically, 62% of Chest pain COVID patients (within the 1st month of the infection) were diagnosed as long COVID while only 32% of patients without Chest pain (within the 1st month of the infection) developed long COVID, which suggests that early chest pain is a good indicator for long COVID. Early joint pain (within the 1st month of the infection) is also a good indicator for long COVID. Namely, 62% of patients with early joint pain developed long COVID while only 30% of patients without early joint pain were eventually diagnosed as long COVID. Age seems to be another important factor for long COVID. We can notice a trend in the proportion of long COVID varying with age. In general, the percentage of COVID infectants who eventually developed long COVID increases along with the aging process. However, the teenage group (9.5-19.0) is an outlier, with the highest risk of 52% among all age groups. A potential explanation for is that the teenage is more socially engaged and they are very likely to be exposed to other risk factors, which suggests that we may need additional actions (compared to other age groups) to protect our young generations. Another interesting discovery is 'Runny Nose' symptom. The heatmap in Fig.6 demonstrates an anti-correlation with the long COVID. In other words, 30% less patients with early runny nose symptom (within the 1st month of the infection) developed long COVID compared to their counterpart without runny nose symptom. Like Joint Pain and Chest Pain, Shortness of breath is another good indicate for long COVID risk. 58% patients with early shortness of breath developed long COVID, which is a lot higher than the patients without these symptoms were diagnosed as long COVID (26%). Chronic kidney disease is our negative example for long COVID. The variable is not captured by our model to relate to long COVID, and it is justified by the heatmap. In fact, the proportion of patients who experiences the symptom and has long COVID only increases by 1% compared to those who do not experience the symptoms and has long COVID. It follows that the proportion of patients who do not have long COVID and experiences this symptom decrease by 1% from those who do not have the symptom. As so, it is unlikely that this variable is related to long COVID. The heatmap shows that variables like "Diarrhea" and "Joint pain" have positive correlation with long COVID. Vaccination has a negative correlation since it gives patient immune power against the disease. More interestingly, "Runny nose" also has a strong negative correlation with long COVID. This might indicate that "Runny nose" is an indicator of an underlying biological process that acts against long COVID.
